# Mucins shed from the laminated layer in cystic echinococcosis are captured by Kupffer cells via the lectin receptor Clec4F

**DOI:** 10.1101/2022.10.06.511139

**Authors:** Anabella A. Barrios, Camila Mouhape, Leonard Schreiber, Linyun Zhang, Juliane Nell, Mariana Suárez-Martins, Geraldine Schlapp, María Noel Meikle, Ana Paula Mulet, Tsui-Ling Hsu, Shie-Liang Hsieh, Gustavo Mourglia-Ettlin, Carlos González, Martina Crispo, Thomas F. E. Barth, Cecilia Casaravilla, Stephen J. Jenkins, Álvaro Díaz

## Abstract

Cystic echinococcosis is caused by the larval stages (hydatids) of cestode parasites belonging to the species cluster *Echinococcus granulosus sensu lato*, with *E. granulosus sensu stricto* being the main infecting species. Hydatids are bladder-like structures that attain large sizes within various internal organs of livestock ungulates and humans. Hydatids are protected by the massive acellular laminated layer (LL), composed mainly by mucins. Parasite growth requires LL turnover, and abundant LL-derived particles are found at infection sites in infected humans, raising the question of how LL materials are dealt with by the hosts. In this article, we show that *E. granulosus sensu stricto* LL mucins injected into mice are taken up by Kupffer cells, the liver macrophages exposed to the vascular space. This uptake is largely dependent on the intact mucin glycans and on Clec4F, a C-type lectin receptor which in rodents is selectively expressed in Kupffer cells. This uptake mechanism operates on mucins injected both in soluble form i.v. and in particulate form i.p. In mice harbouring intraperitoneal infections by the same species, LL mucins were found essentially only at the infection site and in the liver, where they were taken up by Kupffer cells via Clec4F. Therefore, shed LL materials circulate in the host and Kupffer cells can act as a sink for these materials even when the parasite grows in sites other than the liver.

## Introduction

The larval stages of the cestode parasites belonging to the species cluster known as *Echinococcus granulosus sensu lato* (distinguishable only by molecular means) cause cystic echinococcosis in livestock animals including sheep and cattle, as well as in humans (1, 2). Most of cystic echinococcosis worldwide, both in in humans and in livestock species, is caused by the species *E. granulosus sensu stricto* (3). Cystic echinococcosis is characterized by the growth within organ parenchymae of bladder-like parasite structures called hydatids. Each hydatid is bounded by a thin cellular layer, in turn protected by a thick acellular coat called the laminated layer (LL) (4–7). The main component of the LL is a meshwork of mucins, heavily decorated by O-glycans that are rich in galactose (8, 9). Hydatids persist in internal organs of immunocompetent hosts for up to decades, and they usually elicit weak local inflammation (10). The growth of the hydatid (potentially through over 3 orders of magnitude in lineal dimension) requires the shedding of materials from the LL (4–7). In human infections, the abundant shed LL particles (termed SPEGs, for <u>s</u>hed <u>p</u>articles of *<u>E</u>. <u>g</u>ranulosus*) are found in the tissue surrounding the parasite and in the draining lymph nodes (11).

The persistence of larval *E. granulosus* infection, the massive nature of the LL, and sheer abundance of shed particles as detected in human infection call the question of how the host’s body disposes of this foreign debris and avoids the potential inflammation that may result from its build-up. As inflammation is detrimental to the parasite (10), it makes biological sense that LL constituents may be evolutionarily perfected for engaging cellular receptors that will ensure non-inflammatory disposal of the shed materials.

Studies on the responses elicited in mouse dendritic cells (DC) and macrophages by *in vitro*-produced LL particles (pLL, from particulate LL) have suggested an interaction mode independent of conventional receptors and inefficient phagocytosis, at least in the absence of opsonization (7, 12, 13). In broad agreement, none out of 35 human innate carbohydrate-recognition proteins (recombinantly expressed as fusions with IgG Fc) bound the LL mucins in an *in vitro* screening (14). However, in the same experiment, strong binding was observed for mouse Clec4F. This type II, trimeric, cell-surface C-type lectin has affinity for glycans terminated in N-acetylgalactosamine or galactose (Gal), especially when they occur in tandem along the same backbone, as it happens in mucins (15–19). The binding of recombinant Clec4F to the LL mucins was as expected calcium-dependent and resistant to digestion of non-mucin proteins in the parasite preparation. Binding was also observed to a number of synthetic glycans representing some of the main carbohydrate motifs in the *E. granulosus* LL and/or in the LL of the related species *Echinococcus multilocularis* (14).

In rodents, Clec4F is exclusively expressed in Kupffer cells (KC) (15–17), the sessile liver macrophages exposed to the vascular space. This restricted expression means that Clec4F is not expressed in the mouse DC and macrophage sub-types used in the previously mentioned experiments that suggested a receptor-independent interaction mode with LL particles (7, 12, 13). In the ungulate natural hosts of *E. granulosus*, Clec4F appears to be expressed in lymphoid tissues in addition to the liver (16). In humans, Clec4F is not functional as a lectin and is poorly or not expressed (16).

The only demonstrated function of Clec4F (in mice) is the removal from circulation of de-sialylated platelets by KC (20). Although the macrophage galactose-type lectins (MGL) and the asialoglycoprotein receptor (ASGR; reported to be expressed in KC in addition to in hepatocytes) collaborate in this function, Clec4F appears to fulfil an indispensable role (20–22).

KC under basal conditions have a tolerogenic profile as antigen-presenting cells (23). Also, a role of Clec4F in removing de-sialylated platelets (20), conceptually akin to the role of the receptors involved in phagocytosis of apoptotic cells (24), suggests that Clec4F is not wired to inflammatory responses. Thus, Clec4F/KC appears to be an evolutionary option from the parasite’s standpoint for LL debris disposal. However, *in vitro* binding by the soluble Clec4F lectin domain expressed in dimeric form (14) does not predict binding by the native cell-surface trimeric receptor, or internalization by KC. In this article, we analyse *in vivo* uptake of soluble and particulate *E. granulosus sensu stricto* LL materials by KC and the role of Clec4F in this process, in the context of artificial injection of materials as well as experimental infection.

## Materials and Methods

### Parasite materials

The starting material for LL mucin preparations, and the source of protoscoleces for experimental infections, were macroscopically healthy, *E. granulosus sensu stricto* hydatids containing live protoscoleces (i.e. fertile), from natural infections in cattle from Uruguay. Parasite samples were genotyped by amplification and sequencing of mitochondrial cytochrome c oxidase according to (25) and all samples in the study corresponded to genotypes G1 or G3. For LL mucin preparations, hydatid walls (which when arising from bovine hosts have the LL as the overwhelmingly dominant component (4, 5, 9)) were extensively washed with phosphate-buffered saline (PBS), then with 2 M NaCl and finally with water. The walls were then dehydrated by sequential steps with ethanol (95% v/v) followed by acetone, and finely ground using a mortar and pestle, followed by grinding between two polished glass plates (13). The resulting fine powders were rehydrated in endotoxin-free PBS containing 30 mM EDTA (1 ml for every 2 mg dry mass) to extract the calcium inositol hexakisphosphate deposits, thus leaving the mucin meshwork as single major structural component (4). The material was then extensively washed into endotoxin-free PBS. To obtain a fine particulate preparation (pLL), the suspension was filtered through 85- and 23-µm gauze (13). To obtain the soluble preparation (sLL), the suspension was subjected to sonication followed by centrifugation for 1 h at 100,000 g at 4°C. To obtain sLL in which the terminal monosaccharide residues were oxidized, prior to the solubilization step, the material was treated with sodium periodate followed by sodium borohydride (or with sodium borohydride only as mock treatment) (13). For obtaining biotin-tagged sLL, prior to the solubilization step, the material in PBS was incubated overnight with N-hydroxysuccinimido-biotin (NHS-biotin; 10 ng per mg of LL dry mass). Sodium periodate, sodium borohydride and/or NHS-biotin were removed by extensive washing of the particles into endotoxin-free PBS. Destruction of terminal monosaccharides by periodate was verified in terms of abrogation of the binding biotinylated PNA (26), revealed with the help of streptavidin-perodixase, in a dot-blot format. Biotin incorporation by the sLL was checked using streptavidin-peroxidase also in dot-blot format. The dry mass content of LL mucin preparations was estimated as in (13). LL mucin preparations were stored at 4°C in PBS with penicillin-streptomycin during no more than 6 months (pLL) or 12 months (sLL).

### Clec4f^-/-^ mouse line generation

Although Clec4f^-/-^ mice had been previously generated by one of the authors’ group (17), the cost of importing the mice from Taiwan to Uruguay and re-deriving them was high enough that made *de novo* generation of a Clec4f^-/-^ line by CRISPR-Cas9 technology economically convenient. All animal procedures were performed at the Specific Pathogen Free animal facility of the Laboratory Animal Biotechnology Unit of Institut Pasteur de Montevideo. Experimental protocols were approved by the Institutional Animal Ethics Committee (protocol number 007-18), in agreement with National law 18.611 and international animal care guidelines (Guide for the Care and Use of Laboratory Animals) (27) regarding laboratory animal’s protocols. Mice were housed in individually ventilated cages (Tecniplast, Milan, Italy) containing chip bedding (Toplit 6, SAFE, Augy, France), in a controlled environment at 20 ± 1°C with a relative humidity of 40-60%, in a 14/10 h light-dark cycle. Autoclaved food (Labdiet 5K67, PMI Nutrition, IN, US) and autoclaved filtered water were administered *ad libitum*. Cytoplasmic microinjection was performed in C57BL/6J zygotes using a mix of 25 ng/µl Cas9 mRNA (Invitrogen, Carlsbad, CA, US), 15 ng/µl of each sgRNA (2 guides were used) (Synthego, Menlo Park, CA, US), diluted in standard microinjection buffer (5 mM Tris-HCl, pH 7.4, 0.1 mM EDTA). Viable embryos were cultured overnight in M16 (Sigma, St Louis, MO, US) microdrops under embryo tested mineral oil (Sigma), with 5% CO_2_ in air at 37 °C. The next day, 2-cell embryos were transferred into the oviduct of B6D2F1 0.5 days postcoitum pseudopregnant females (20 embryos/female in average). Standard surgical procedures established in our animal facility were followed (28). For surgery, recipient females were anesthetized with a mixture of ketamine (110 mg/kg, Ketonal50, Richmond Veterinaria S.A., Buenos Aires, Argentina) and xylazine (13 mg/kg, Seton 2%; Calier, Montevideo, Uruguay). Tolfenamic acid was administered subcutaneously (1 mg/kg, Tolfedine, Vetoquinol, Madrid, Spain) in order to provide analgesia and anti-inflammatory effects (29). Pregnancy diagnosis was determined by visual inspection by an experienced animal caretaker two weeks after embryo transfer, and litter size was recorded on day 7 after birth. Pups were tail-biopsied and genotyped 21 days after birth, and mutant animals were maintained as founders. Gene editing efficiency was 53.8% (7 KO/13 born pups). Genetic modification was confirmed by Sanger sequencing. The Clec4f^-/-^ line was established after producing F1, F2, and finally homozygous F3 animals that were used for experiments. Primers used for amplification of the genomic locus were 5’-TCTTTATGATCGCACCCACA-3’, 5’-TCCATTCTCGAGAGCCATCT-3’. F1 and F2 heterozygous mice were selected using a heteroduplex mobility assay on a 10% polyacrylamide gel. In F2, WT animals were differentiated from homozygous KO animals by Sanger sequencing.

### Mouse experiments

WT BALB/c mice were obtained from DILAVE (MGAP, Uruguay) and kept at the Instituto de Higiene animal house. Female WT (C57BL/6J) or Clec4f^-/-^ mice were bred at the Institut Pasteur de Montevideo, co-housed (so as to equilibrate their microbiota) at the Instituto de Higiene starting at 5 weeks of age for at least 3 weeks, and used between 8 and 14 weeks of age. sLL (100 or 200 μg dry mass per mouse as indicated in each case, dissolved in 100 μl of PBS) or vehicle only was injected into the tail vein. pLL (450 μg dry mass per mouse, suspended in 200 μl of PBS) or vehicle only, were injected i.p. Mice were infected by i.p. injection of 2000 (BALB/c) or 2500 (C57BL/6) protoscoleces (viability ≥ 95%) suspended in 200 μl of PBS; control mice were injected with vehicle only. Mice were euthanized using isoflurane. The procedures used were approved by the Comisión Honoraria de Experimentación Animal (CHEA, Universidad de la República, Uruguay: protocols 101900-001017-16, 101900-000994-16, 101900-000649-17, 101900-000441-16).

### Preparation of liver non-parenchymal fraction for flow cytometry

Following perfusion through the inferior vena cava with 10 mL of PBS, livers were placed in RPMI and finely chopped using a razor blade for approximately 2 min. The tissue was then digested in 5 ml per g of enzyme mix (RPMI containing 0.4 U/ml Liberase [Sigma] and 80 U/ml DNase [Sigma]), for 25 min at 37◦C, in an orbital shaker at 250 rpm with additional manual shaking every 5 min. Digests were poured through a 100 µm strainer and centrifuged at 300 g for 5 min; the cell pellets were re-suspended in 25 ml of RPMI and again centrifuged at 300 g. The pellets were suspended in 2 ml of RBC lysis buffer (Sigma), and after 2 min, 2 ml of FACS buffer (PBS supplemented with 0.5% BSA and 2 mM EDTA) was added. After another centrifugation step (300 g, 5 min), cells were re-suspended in 3 ml of FACS buffer, passed through a 40 µm strainer, and counted using the Nexcelom Cellometer K2 with the AOPI stain for cell viability. Cells and buffer used with the cells were kept at 4°C, except for the digestion step.

### Flow cytometry

Five hundred thousand live cells where incubated with the viability probe for 10 min at room temperature, then with 0.025 µg of anti-CD16/32 (Biolegend) in 10% v/v normal rat serum (Unidad de Reactivos y Biomodelos de Experimentación, Facultad de Medicina, Universidad de la República). Cells were then surface-stained, fixed in 1% w/v PFA for 20 min at 4°C, washed, and permeabilized overnight with permeabilization buffer (Thermo) for intracellular staining. The details of the viability probes, antibodies and lectin probes are given in Supplementary Table 1. Finally, cells were washed, spun at 350 g for 5 min and re-suspended in FACs buffer. The data were acquired in a FACS Canto II Cytometer (BD), at the Instituto de Higiene, Universidad de la República, and then analysed using the FlowJo software package. The relative contribution of each liver non-parenchymal cell type under study to the uptake of LL materials was estimated by multiplying the frequency of cells positive for the LL probe and belonging to the particular cell type (within all live single cells) by their arithmetic mean fluorescence intensity for the probe and then dividing this figure over the summation of the analogous products for the 3 cell types analysed.

### Histology

Tissue pieces (approximately 0.5 mm^3^) were fixed in buffered formalin. Liver samples corresponded to animals subjected to liver perfusion with PBS (as the materials were also analyzed by flow cytometry). Paraffin-embedded tissue sections (2 µm) were probed for either chromogenic or two-color fluorescent detection. For chromogenic detection, sections were probed with biotinylated monoclonal antibody E492 (3 µg/mL; steamer pH 6.1) or biotinylated PNA (Vector Laboratories, Newark, California, United States, B-1075; dilution 1:100; steamer pH 9.0) using the avidin-biotin-complex method (30), followed by Dako REAL™ Detection System, Alkaline Phosphatase/RED, Rabbit/Mouse (Agilent, Santa Clara, California, USA), and counterstained using haemalum. For fluorescent detection, sections were probed with biotinylated monoclonal antibody E492 (3 µg/mL; steamer pH 6.1) alongside a monoclonal antibody either against Clec4F (clone 370901, rat IgG2a, Thermo Fisher Scientific, Waltham, Massachusetts, USA; 1:50) or against F4/80 (clone BM8; Invitrogen, Waltham, Massachusetts, USA; 1:100). The fluorochrome used for E492 labelling was Streptavidin Alexa Fluor 488 (Thermo Fisher Scientific, Waltham, Massachusetts, USA; 1:1600). For Clec4F and F4/80, we used polyclonal goat IgG anti rat IgG coupled to biotin (Dianova, Hamburg, Germany; 1:100), followed by Streptavidin Alexa Flour 546 (Thermo Fisher Scientific, Waltham, Massachusetts, USA; 1:1000). The counterstain was performed with DAPI (4′,6-diamidino-2-phenylindole). Negative controls were performed by omitting the primary antibody. Slides were evaluated in an Axiophot microscope (Oberkochen, Germany) coupled to a CCD camera.

### ELISA

ELISA wells were coated with sLL at 10 µg/mL (total dry mass) in Tris-buffered saline (TBS) overnight at 4 °C, and blocked using TBS containing 0.05% w/v Tween 20, 1% w/v bovine serum albumin and 0.5% v/v Carbo-Free^TM^ blocking solution (Vector Laboratories). Wells were then probed with biotinylated E492 antibody (31), biotinylated peanut agglutinin (PNA; Vector laboratories) or recombinant soluble Clec4F (expressed as a fusion with the F_c_ fragment of human IgG_1_ (32)). The biotinylated PNA or E492 antibody were detected using streptavidin coupled to peroxidase (ThermoFisher Scientific) and the recombinant Clec4F was detected using a goat anti-human IgG coupled to peroxidase (AbCam, Cambridge, UK). For competition ELISA, recombinant Clec4F at 80 ng/mL was mixed with variable concentrations of either (biotinylated) PNA or E492 antibody as competitors, and bound Clec4F detected as previously described.

### Particle immunofluorescence

pLL particles were incubated with biotinylated E492 antibody (0.6 µg/mL), biotinylated PNA (1:1500) or recombinant soluble Clec4F (7.5 µg/mL), followed by streptavidin-Cyanine-3 (1:500; Amersham Bioscience) or by goat anti-human IgG coupled to FITC (1:100, Sigma), as appropriate. Control stains were included using human IgG_1_ instead of recombinant Clec4F as well omitting the primary staining reagents.

### Statistical analyses

As the underlying data do not satisfy the criteria for parametric statistics, rank-based nonparametric statistical methods, were used throughout. When the numbers of mice in each treatment-repeat experiment combination were equal, the Mack-Skillings exact test for a two-way layout was used (33). When the data did not fulfil the above condition, data from the different repeat experiments were pooled and the modified Wilcoxon-Mann-Whitney test (only two experimental groups) (53) or the Kruskal-Wallis test (more than two experimental groups) were used. When the Mack-Skillings or the Kruskal-Wallis tests resulted in a statistic with a p-value less than 0.05, the test was followed by the post-hoc multiple comparison test described by Conover (55) and the Benjamini-Hochberg correction for controlling the false discovery rate (56). Throughout the paper, the symbols *, ** and *** represent p-values less than 0.05, 0.01 and 0.001 respectively.

## Results

### Soluble LL mucins are taken up by Kupffer cells in vivo in a manner dependent on the mucin glycans

We first sought to determine whether KC take up soluble LL mucins (sLL) *in vivo* and whether this is dependent on the mucin glycans. We thus treated the LL mucins either with sodium periodate to oxidize the glycans or applied a mock treatment, tagged both preparations with equivalent amounts of biotin (Fig. S1), and injected both preparations i.v. in mice. Twenty minutes later, we quantitated the biotin tag in permeabilized non-parenchymal liver cells using streptavidin conjugated to PE and flow cytometry. KC were identified using our previously validated gating strategy (Fig. S2) (34). A significant signal was detected in KC, and the signal was markedly higher for mock-treated than for periodate-treated sLL (Fig. 1). No significant biotin signal was detected in either liver sinusoidal epithelial cells (LSECs) or in non-KC CD11b^+^ cells, a broad definition that encompasses any mononuclear phagocytes other than KC present in the liver (Fig. 1 a, b, and S2).

**Fig. 1.**
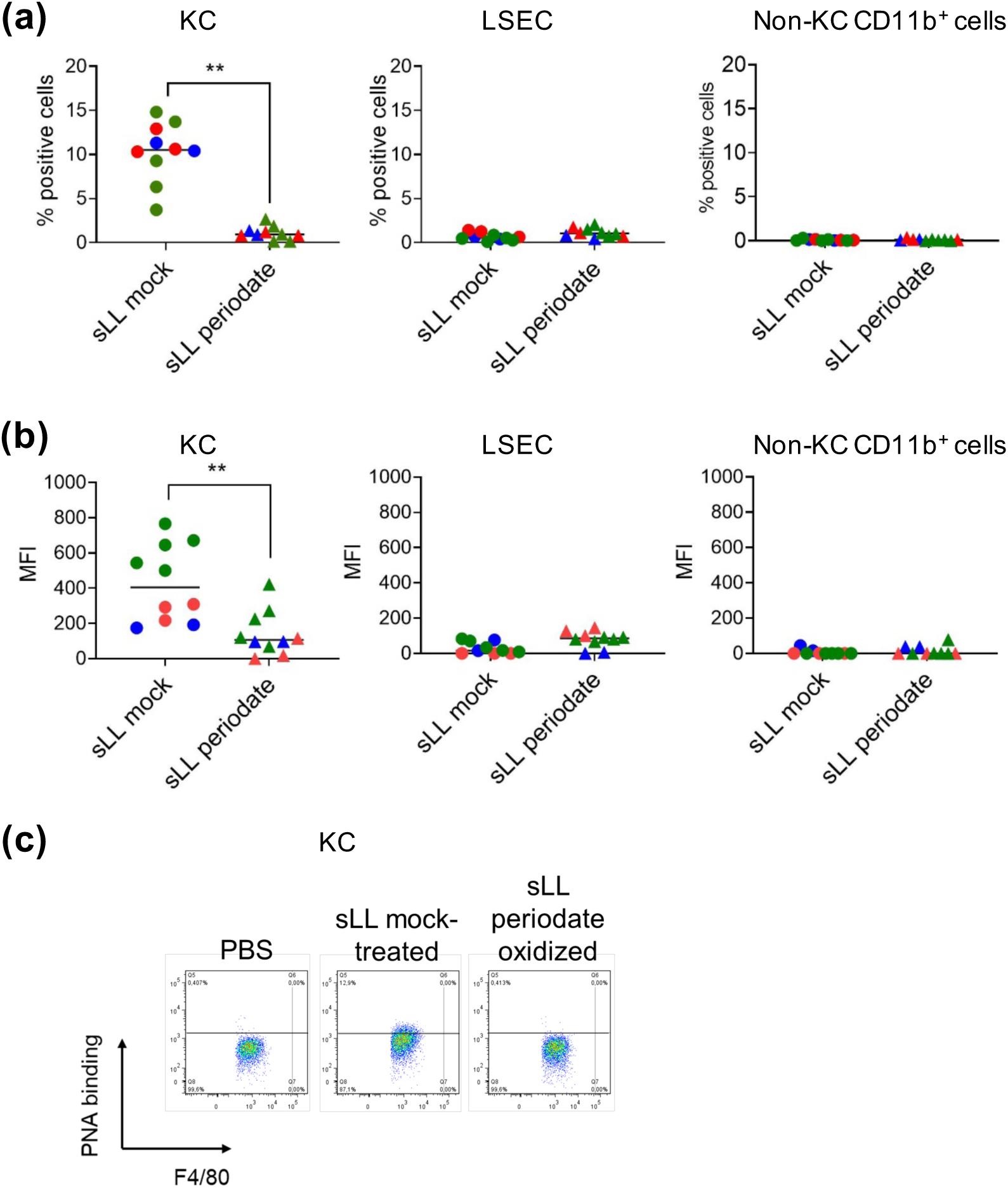
Soluble LL mucins are taken up selectively by KC, dependent on mucin glycans. C57BL/6 mice were injected i.v. with 100 μg dry mass of solubilized, biotin-derivatized LL mucins (sLL), either treated with periodate to oxidize their terminal monosaccharide residues or mock-treated. Twenty minutes later, mice were sacrificed and sLL uptake was quantitated in permeabilized liver non parenchymal cells using streptavidin-PE. Data are shown for the cell types that are relevant in the context of the article, namely KC, LSEC and non-KC CD11b^+^ cells, in terms of percentages of positive cells **(a)** and geometric means of fluorescence intensity **(b)**. Representative dot-plots for KC are shown in **(c)**. The signal threshold for each cell type was established using cells from mice injected with vehicle only and similarly stained; the fluorescence intensities data had the median values corresponding to vehicle-injected mice subtracted for plotting. Results are pooled from 3 independent experiments (represented in different colours); individual mice and their overall median values are shown. The statistical analyses were carried out on the crude data, by the Mack-Skillings test.

### The plant lectin PNA and the E492 monoclonal antibody allow detection of laminated layer mucins captured by liver cells in vivo

Except in experiments using periodate-treated LL materials, the detection of LL materials by appropriate carbohydrate-binding probes was preferable to prior labelling with biotin. Among other advantages, such probes would allow the detection of endogenous LL mucins in infected animals. One candidate probe was the plant lectin PNA, which binds Galβ1-3-GalNAcα1-peptide, i.e. the non-decorated mucin O-glycan core 1 or T-antigen (35). This motif is highly represented in the *E. granulosus* LL, and accordingly, PNA binds the LL mucins in histological sections, particle lectin fluorescence, and Western blotting (13, 26). The second candidate probe was the monoclonal antibody E492, raised against *E. granulosus* protoscoleces (36). This antibody binds to LL mucins from *E. multilocularis* (37) and to synthetic glycans that include the P_1_ blood antigen motif (Galα1-4Galβ1-4GlcNAc), highly represented among *E. granulosus* LL mucin glycans (8, 9). A third potential probe was recombinant Clec4F (14). We verified that these potential probes indeed bind to sLL coated on ELISA plates as well as to LL particles in suspension (Fig. 2 a, b).

**Fig. 2.**
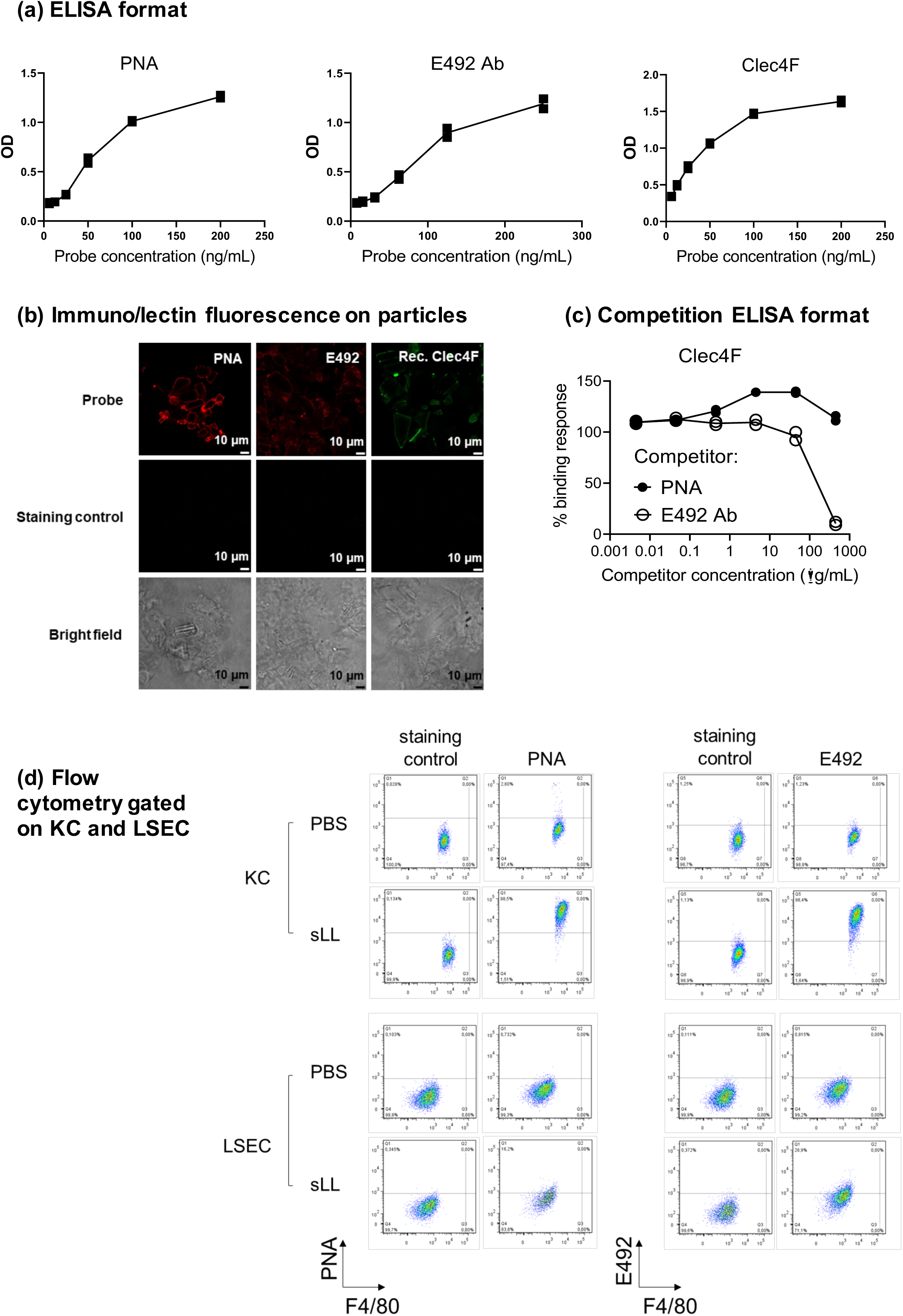
PNA and the E492 antibody are suitable probes for LL mucins captured by mouse KC. The binding of PNA, the E492 antibody and soluble recombinant Clec4F to LL mucins coated onto ELISA plate wells **(a)** or present in LL particles in suspension **(b)** was studied. The capacity of PNA and the E492 antibody to compete with the binding of recombinant Clec4F to LL mucins was analysed in a competition ELISA format **(c)**; results are expressed in relation to the response in the absence of competitor. In (a) and (c), analytical duplicates are shown. WT C57BL/6 mice were injected i.v. either with PBS only or with solubilized LL mucins (200 µg/mouse) and 22 h later KC and LSEC were analysed by flow cytometry using either PNA or the E492 antibody as probes, after cell permeabilization (**d**). Staining controls shown in (b) and (d) correspond to samples stained in the same way as the positive ones except that the primary probe for LL material was omitted, and the analogous controls in (a) gave no signal.

We took advantage of the defined specificities of PNA and the E492 antibody to gain insight into the carbohydrate motifs bound by Clec4F in the LL mucins. The binding of recombinant Clec4F to the solubilized and plate-bound LL mucins was not competed by PNA, but it was strongly competed by the E492 antibody (Fig. 2 c). Therefore, most of the binding of Clec4F to the native LL mucins appears to rely on recognition of the P_1_ motif, whereas recognition of the T-antigen is not significant. This in turn suggests that in the LL mucins the T-antigen may not be exposed and/or be spaced inadequately for Clec4F binding, as recombinant Clec4F does bind the T-antigen in glycan arrays (16–18).

*In vivo* probes should show minimal reactivity with host tissues; for our purposes, lack of reactivity with the populations of mouse non-parenchymal liver cells under study was the most important consideration. Neither PNA nor the E492 antibody showed significant binding in flow cytometry to KC or LSEC from control mice (Fig. S3). PNA bound to a very minor fraction of non-KC CD11b^+^ cells, whereas the antibody did not bind. PNA, but not the E492 antibody, bound to a major sub-population of CD45^+^ CD11b^-^ F4/80^-^cells, which were not of particular interest in our study. Recombinant Clec4F bound to cells in all 4 cell gates defined, most strongly to LSEC and KC (Fig. S3). This probably reflects the fairly broad specificity of Clec4F for glycans terminated in Gal/GalNAc (16–18), and ruled out this lectin as a probe for our purposes.

Finally, to determine whether uptake of sLL could be measured *in vivo* using PNA or the E492 antibody, we injected wild-type (WT) C57BL/6J mice i.v. with sLL and subsequently stained isolated hepatic non-parenchymal cells intracellularly with either probe. As expected, no staining was observed in cells from mice injected with PBS only, whereas KC, and to a small extent also LSEC, from mice given sLL stained intracellularly with either PNA or E492 antibody (Fig. 2 d). Omitting the cell permeabilization step caused the PNA signal in KC and LSEC of sLL-injected mice to be reduced to negligible levels (data not shown), ruling out the remote possibility that injection of sLL induces the expression of ligands for the probe in the glycocalix of the cells under study. Both PNA and the E492 antibody appeared to allow more sensitive detection than prior labelling with biotin (Fig. 1), so that captured sLL was detected in almost all KC and also in some LSEC.

In summary both PNA and the E492 antibody are useful probes for LL mucins captured by mouse non-parenchymal liver cells.

### Soluble LL mucins are taken up by Kupffer cells in vivo in a manner dependent on Clec4F

We first used CRISPR/Cas9 to generate a new Clec4f ^-/-^ line on the C57BL/6J background. These Clec4f ^-/-^ mice have 8-bp deletion expected to result in a polypeptide that is truncated near the N-terminus (Fig. 3, a, b), and effectively lack Clec4F protein expression by flow cytometry (Fig. 3 c). Hence, our line appears similar to Clec4f gene-deficient mice that have already been described (17, 20).

**Fig. 3.**
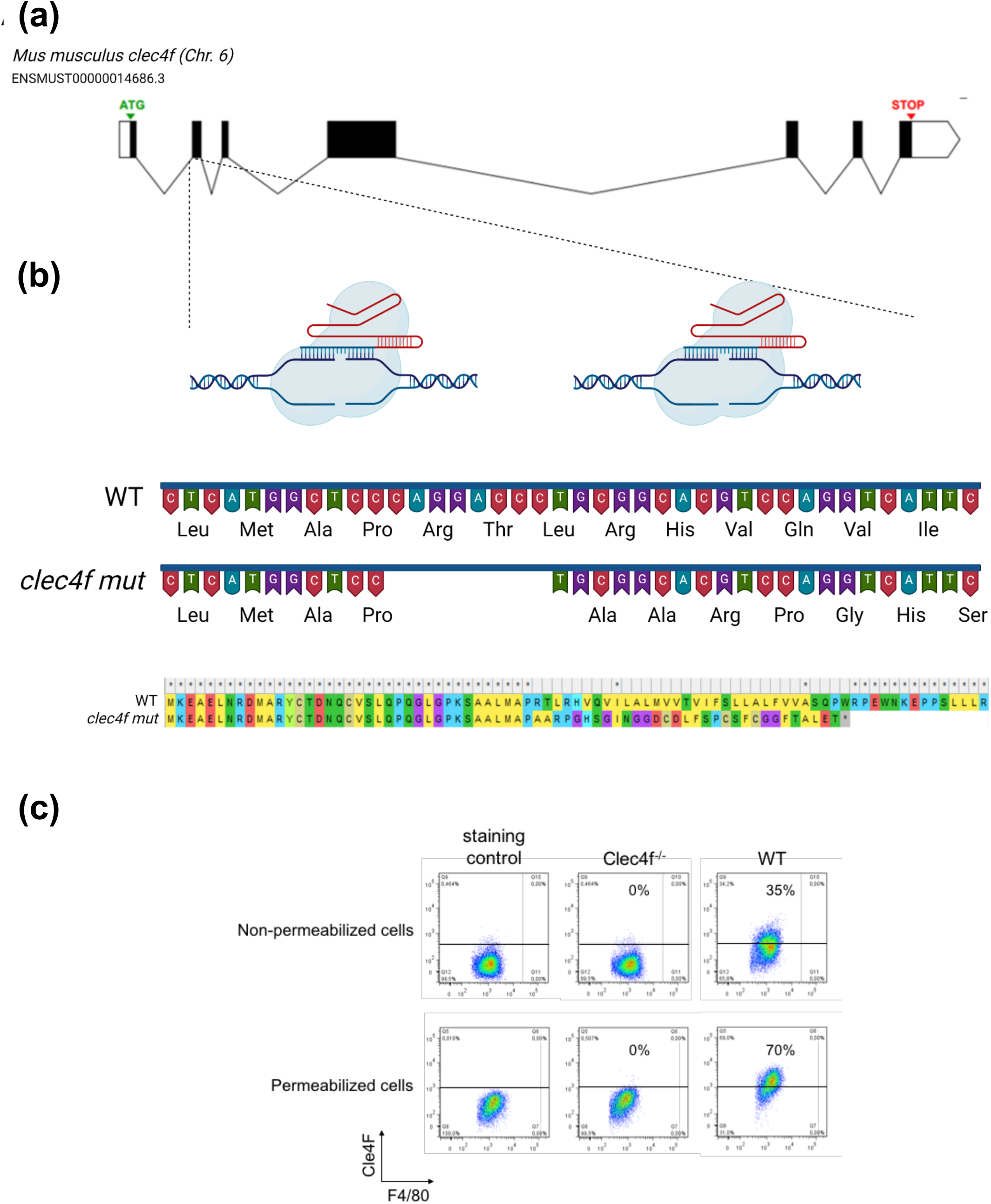
Generation of Clec4f gene-deficient mice. (a) Schematic structure of the mouse *Clec4f* gene. Coding exons appear as solid black bars and non-coding regions as white bars. Introns are represented as lines. **(b)** The target sequences for CRISPR/Cas9, 5’-TGGACGTGCCGCAGGGTCCT-3’ and 5’-GCTTCTCCCCTGCAGGACTG-3’, are located in the second exon and first intron/second exon junction of the gene respectively. An 8 bp deletion corresponding to nucleotides 5’-CAGGACCC-3’ was identified by tail biopsy. The frameshift mutation produces a protein 480 aa shorter. **(c)** The liver non-parenchymal fraction was prepared from Clec4f^-/-^ and WT mice and analyzed for Clec4F expression by flow cytometry without or with cellular permeabilization. KC were defined as shown in Figure S2. The staining controls contain all the fluorophore-coupled antibodies but lack the primary anti-Clec4F antibody.

We next compared WT and Clec4f^-/-^ mice in terms of the *in vivo* uptake of sLL by liver non-parenchymal cells. For this purpose, we injected untagged sLL, either untreated or subjected to the mock treatment used in the previous experiments that involved periodate oxidation. Uptake of sLL by KC was strongly dependent on Clec4F (Fig. 4 a - c), particularly when assessed in terms of mean fluorescence intensity. The low-level uptake by LSEC observed was as expected independent of Clec4F; in fact, for untreated sLL, uptake by LSEC was slightly enhanced in the absence of Clec4F, possibly as a result of the decreased capture by KC. No uptake by non-KC CD11b^+^ cells was detected (Fig. 4 a, b and Fig. S4). Through a calculation that takes into account the numbers of each cell type capturing sLL and the intensity of the signal indicating intracellular sLL, KC were estimated to be responsible for the overwhelming majority of sLL uptake by non-parenchymal liver cells in WT mice (Fig. 4 d). However, in the absence of Clec4F, LSEC played a significantly, if slightly, more important part in the uptake of sLL than in the presence of the receptor. Differences in the uptake of untreated *vs* mock-treated material were not observed, except in the low-level uptake by LSEC in Clec4f^-/-^ mice (Fig. 4 a). For consistency with the results shown in Figure 1, we chose to use mock-treated material in the following experiments.

**Fig. 4.**
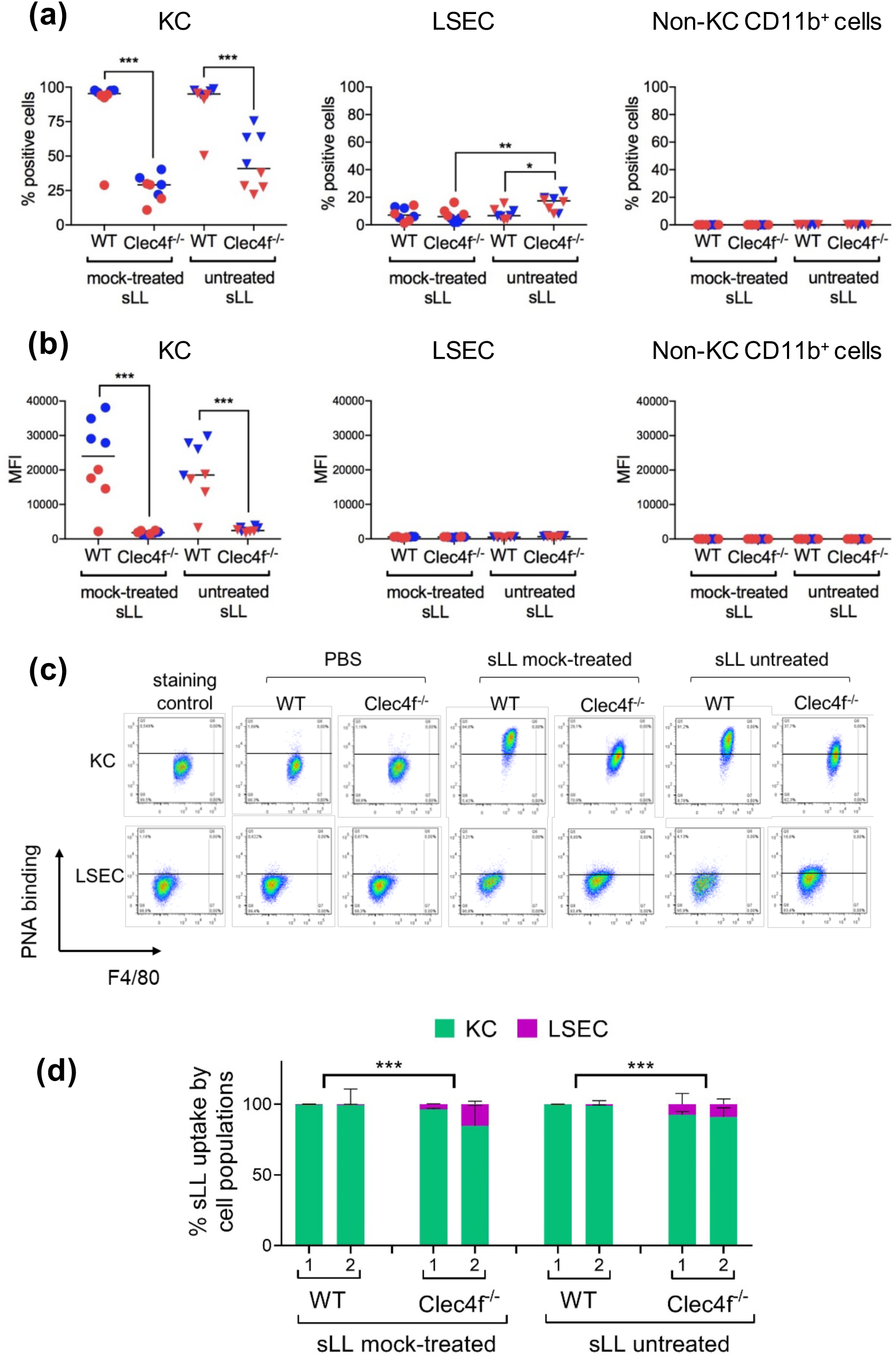
Fast uptake of soluble LL mucins by Kupffer cells depends largely on Clec4F. WT and Clec4f^-/-.^ mice were injected i.v. with 200 ug dry mass of sLL, either untreated or mock-treated as in Figure 1, or vehicle only (PBS). Twenty min later, mice were sacrificed and sLL uptake was quantitated in non-parenchymal liver cell populations by flow cytometry in permeabilized cells using PNA. Results are shown for KC, LSEC and non-KC CD11b^+^ cells, in terms of percentages of positive cells **(a)** and mean fluorescence intensities **(b)**, and representative flow cytometry plots are shown for KC and LSEC **(c)**. An estimation of the fraction of sLL taken up by each of the cell populations under study, calculated as explained in the Materials and Methods section, is shown in **(d)**; the statistics shown compare the fractions taken up by KC. The positive cell threshold was established using cells from the mice injected with PBS only. The data arise from two independent experiments, represented in blue and red in (a-c) and indicated as 1, 2 in (d). In (a-c), individual mice and their median values, and in (d) only medians, are shown. The statistical analyses were carried out by the Mack-Skillings test.

It can be expected that in the absence of Clec4F, other mechanisms of cellular uptake of sLL may become apparent, e.g. macropinocytosis or uptake dependent on natural antibodies or complement (7). Such Clec4F-independent uptake of sLL was hinted by the previous observations in relation to LSEC, as well as by the 25 – 60 % KC in Clec4f^-/-^ mice positive for PNA at low levels (Fig. 4 a - c). As the Clec4F-independent mechanisms might become dominant at longer analysis times, we carried out a similar experiment with a 22 h analysis time. Uptake of sLL was still dominated by KC among the cell types studied, and this uptake was again strongly dependent on Clec4F (Fig. 5 a –d). In the absence of Clec4F, LSEC made a significantly larger contribution to sLL uptake than in the presence of the receptor, but this contribution was still minimal (Fig. 5 d). Non-KC CD11b^+^ cells did not take up material detectably (Fig. 5 a – d and S4). Cell surface expression of Clec4F in KC was not altered by sLL injection (Fig. S5).

**Fig. 5.**
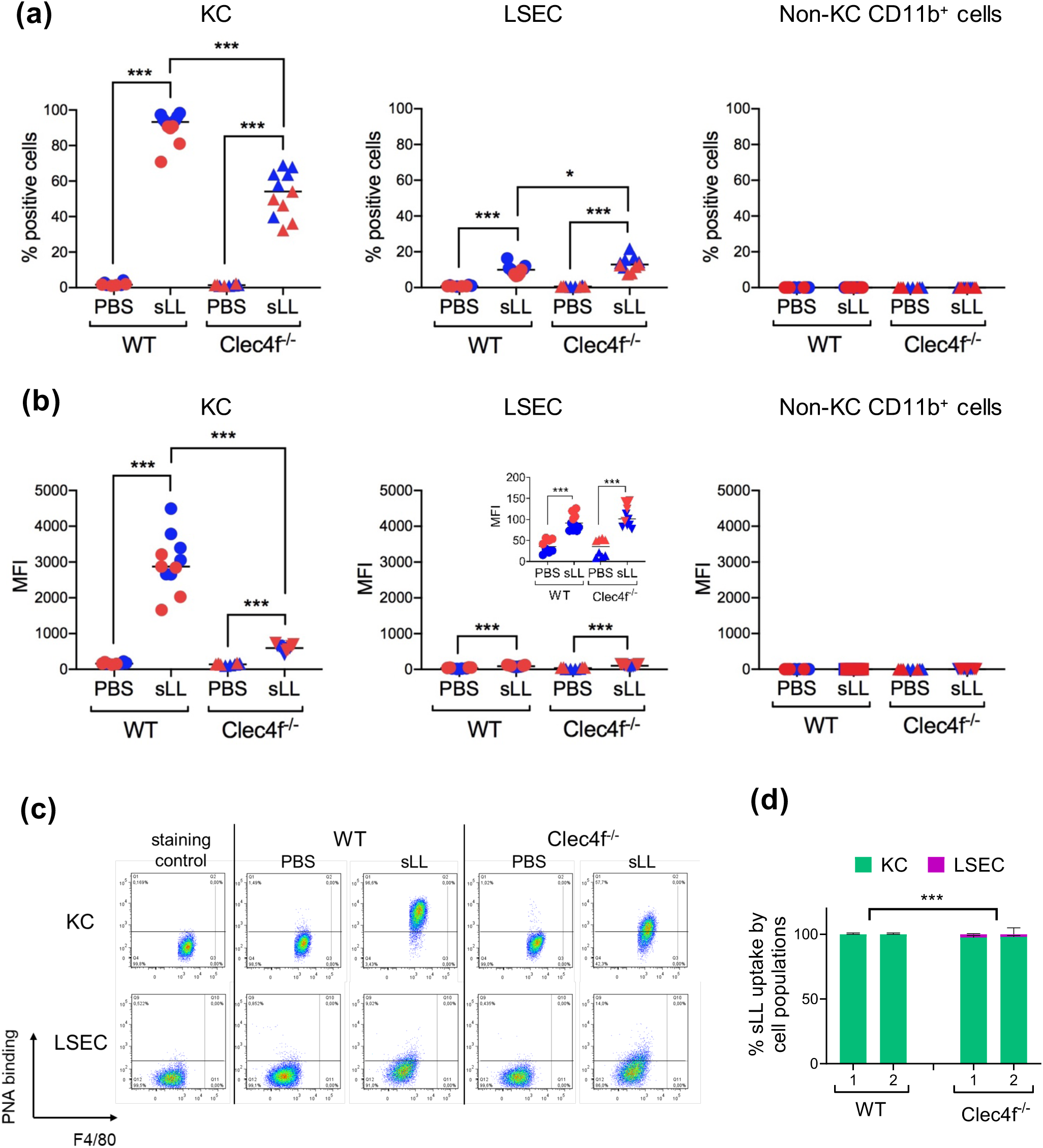
Clec4F is important for the capture of soluble LL mucins by Kupffer cells even after allowing a long uptake time. WT and Clec4f^-/-^ C57BL/6 mice were injected i.v. with 200 µg dry mass per mouse of sLL (mock-treated as in Fig. 1) or PBS only as a control. Twenty-two h later, mice were sacrificed and sLL uptake was quantitated in liver non-parenchymal cells by flow cytometry using PNA after cell permeabilization. Results are shown for KC, LSEC and non-KC CD11b^+^ cells, in terms of percentages of positive cells **(a)** and mean fluorescence intensities **(b)**, and representative flow plots for KC and LSEC are shown in **(c)**. An estimation of the fraction of sLL taken up by each of the cell populations under study, calculated as explained in the Materials and Methods section, is shown in **(d);** the statistics shown compare the fractions taken up by KC. The positive cell threshold was established using staining controls (in which only biotinylated PNA was omitted). The PNA signal in non-permeabilized cells was negligible. The data arise from two independent experiments, represented in blue and red in (a-c) and indicated as 1, 2 in (d). In (a-c), individual mice and their median values, and in (d) only medians, are shown. The statistical analyses were carried out by the Mack-Skillings test (a-c) and the modified Wilcoxon-Mann-Whitney tests (d).

In summary, soluble circulating LL mucins in naïve mice are taken up mainly by KC among liver non-parenchymal cells and mostly in Clec4F-dependent fashion.

### LL mucins injected in particulate form are taken up by Kupffer cells in vivo in Clec4F-dependent manner

Although the release of soluble mucins from the LL during cystic echinococcosis is a possibility (7), only particles have been documented to be shed from the LL in human infections (7, 11). It is not known if these particles circulate systemically and/or give rise to soluble materials that circulate. It was therefore important to determine whether Clec4F participates in the uptake by KC of LL mucins released in a particulate presentation into the internal milieu and at an anatomical site distant from the liver. We thus injected i.p. particulate LL mucins (pLL) (13), and analysed the liver non-parenchymal fraction 22 h later. LL material was indeed detected by flow cytometry in KC (Fig. 6 a – c and S6 a - c). Uptake by KC was dependent on Clec4F, although there was a considerable degree of Clec4F-independent uptake as well. In addition to low-level uptake by LSEC, in this experimental system, non-KC CD11b^+^ cells were observed to take up the LL material. Although non-KC CD11b^+^ cells may include recruited monocytes/macrophages, the frequency of this population (within all non-parenchymal cells) did not increase as a result of pLL injection (Fig. S7). As expected, uptake by LSEC and by non-KC CD11b^+^ cells was independent of Clec4F and in fact tended to be enhanced in the absence of the receptor (Fig. 6 a – c and S6 a - c). Indeed, whereas in WT mice the overwhelming majority of the LL material uptake by liver non-parenchymal cells was carried out by KC, in Clec4f^-/-^ mice the two non-KC cell types studied taken together played a significantly larger role than in WT mice (Fig. 6 d and S6 d). The cell surface expression of Clec4F in KC was not altered by injection of pLL (Fig. S8).

**Fig. 6.**
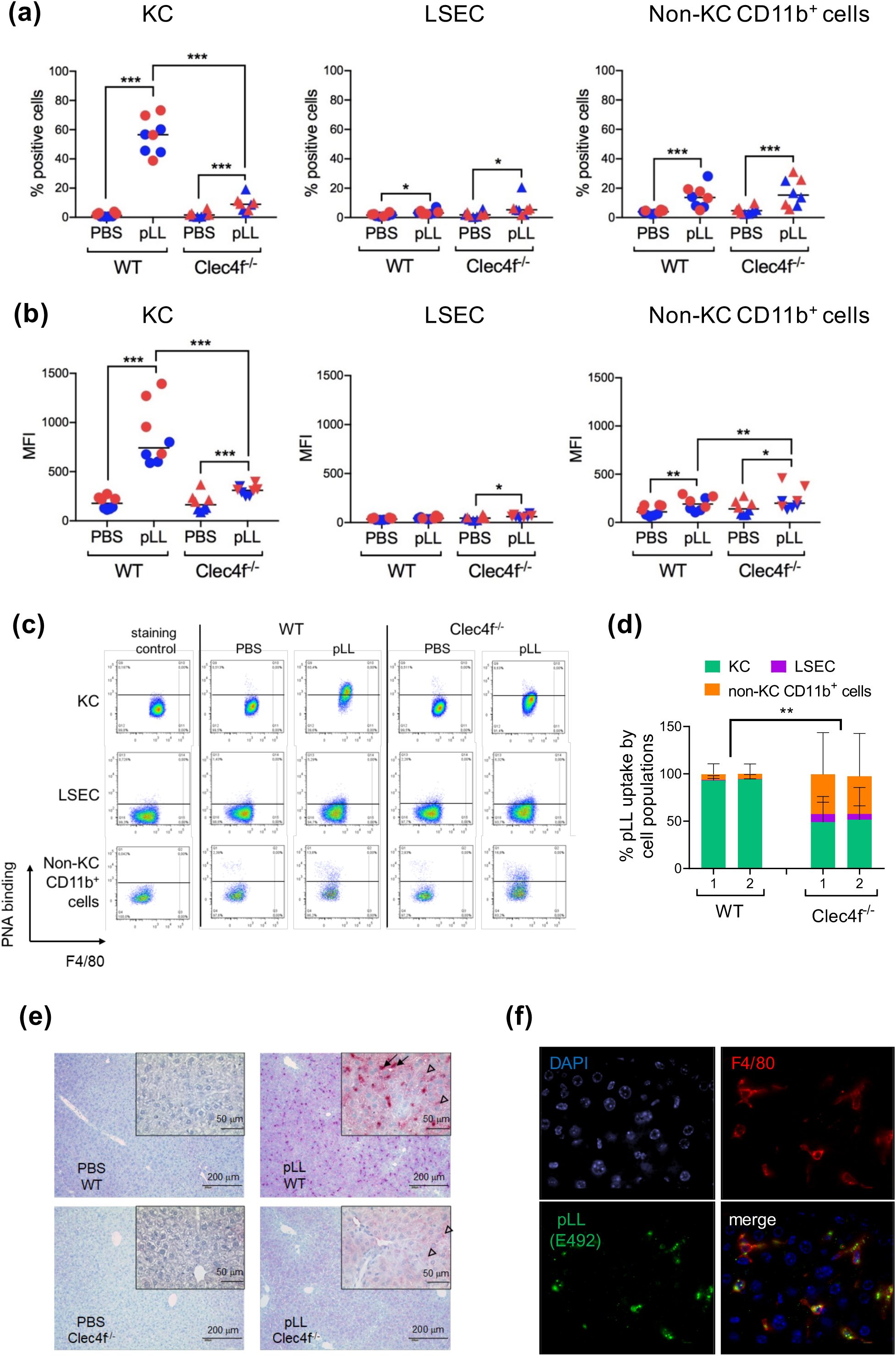
The uptake by Kupffer cells of LL mucins injected in particulate presentation depends largely on Clec4F. WT and Clec4f^-/-^ C57BL/6 mice were injected i.p. with 450 µg dry mass of pLL or PBS only as a control. Twenty-two h later, mice were sacrificed and pLL uptake was quantitated in liver non-parenchymal cells by flow cytometry using PNA after cell permeabilisation. Results are shown for KC, LSEC and non-KC CD11b^+^ cells, in terms of percentages of positive cells **(a)** and mean fluorescence intensities **(b)**, and representative dot-plots for the three cell types are shown in **(c)**. An estimation of the fraction of sLL taken up by each of the cell populations under study, calculated as explained in the Materials and Methods section, is shown in **(d)**; the statistics shown compare the fractions taken up by KC. The positive cell threshold was established using staining controls. The data arise from two independent experiments, represented in blue and red in (a-c) and indicated as 1, 2 in (d). In (a-c), individual mice and their median values, and in (d) only medians, are shown. The statistical analyses were carried out by the Mack-Skillings test. The analogous flow cytometry results obtained using the E492 antibody for detection are shown in Fig. S6. In addition, pLL was detected in liver sections, using the E492 antibody, by chromogenic histology (**e**). Black arrows indicate KC-like cells staining strongly for LL material and empty arrowheads indicate hepatocytes staining weakly for LL material. The localization of LL materials in KC from infected mice was confirmed by fluorescence histology, using the E492 antibody as probe and F4/80 as KC marker **(f)** (original magnification x630).

The mostly Clec4F-dependent uptake of the LL material was confirmed by histology, using the E492 antibody as a probe. Strong staining was observed in KC-like cells of WT animals, whereas only minimal staining was observed in the KC-like cells of Clec4f^-/-^ animals (Fig. 6 e). These cells were formally identified as KC by dual-color fluorescence using F4/80 as KC marker and antibody E492 as probe for pLL (Fig. 6 f). Animals injected with pLL also showed weak staining in hepatocytes (a cell type not analysed by flow cytometry), and this was independent of Clec4F (Fig. 6 e). As hepatocytes are unlikely to phagocytose large particles, this observation suggests the particles injected release small particles and/or soluble mucin materials *in vivo* that preferentially circulate systemically.

In summary, LL mucins present within tissues in particulate form can generate materials (particulate and/or soluble) that circulate systemically and that are captured by KC via Clec4F.

### In experimental E. granulosus infection, LL mucins circulate and are captured selectively by Kupffer cells, with the participation of Clec4F

We next wished to determine if Clec4F was relevant in the infection setting. The most widely used experimental infection model for *E. granulosus sensu lato* uses intraperitoneal injection with protoscoleces, parasite stages that usually develop into adult worms but are capable of reverse development into hydatids (1, 38). In this model, hydatids develop free or within host collagen capsules in the peritoneal cavity (39). KC uptake of LL materials in this model would require that such materials reach systemic circulation, a possibility for which there was no evidence, in experimental or natural infections. We first analysed infected mice of the BALB/c strain, in which experimental infection reaches higher parasite burdens than in the C57BL/6 strain (40). Indeed, the BALB/c mice analysed had a median of 25 mL of total parasite volume (Fig. S9 a), an extremely high parasite burden. In these mice, using the E492 antibody as probe, we detected LL materials by histology in peritoneal cavity cells (i.e. the site of infection) and the liver, but not in kidneys, intestine or lungs (Fig. 7 a). In addition, in some of the infected mice, occasional positive cells were observed in the spleen, and less frequently in the mesenteric lymph nodes (Fig. 7 a). The use of PNA as probe yielded similar results, but it was complicated by the expected signal in the lymphoid organs (41) unrelated to LL material, i.e. detectable in non-infected mice (data not shown).

**Fig. 7.**
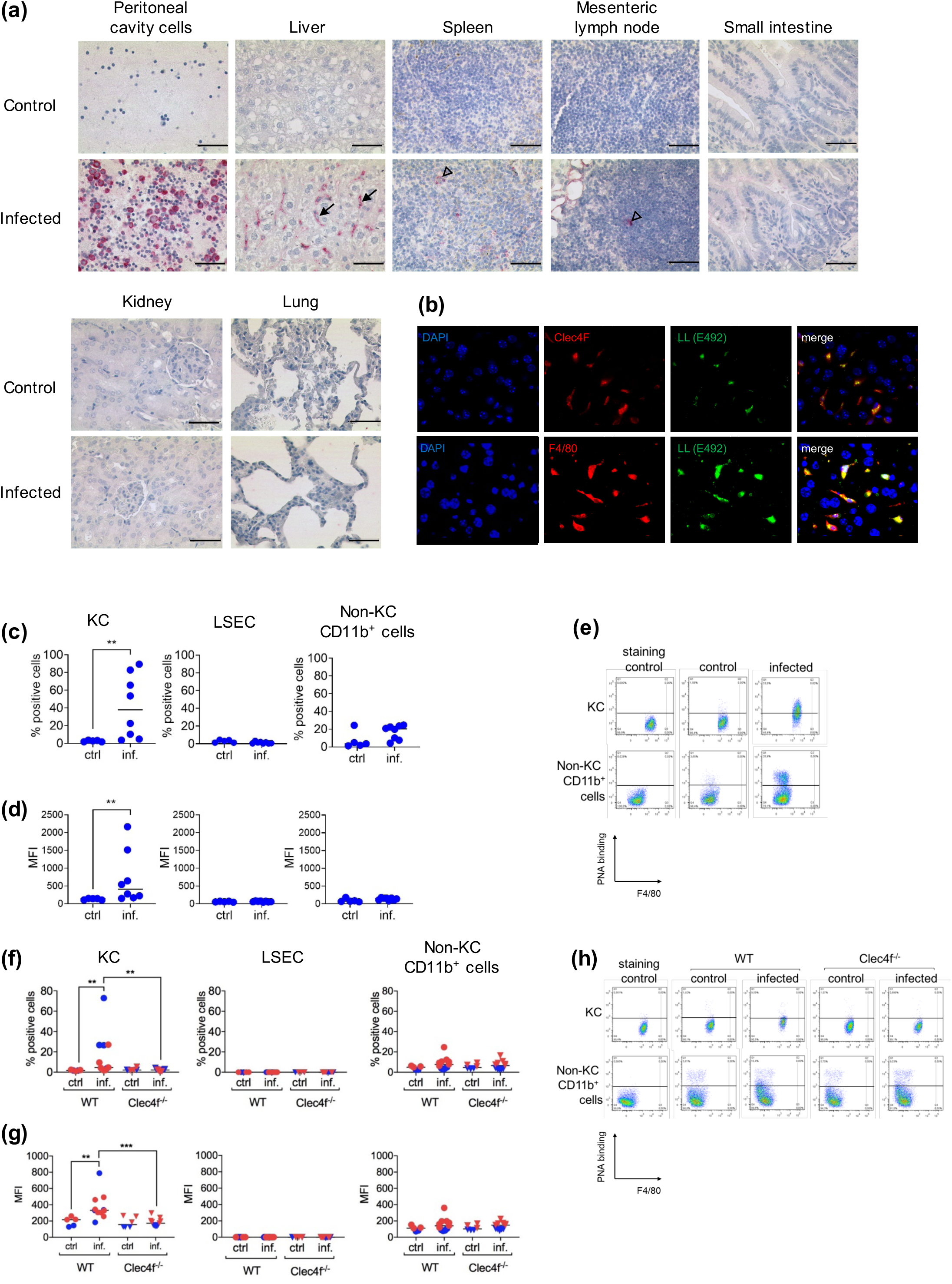
LL materials circulate systemically in infected mice and are captured selectively by Kupffer cells via Clec4F. BALB/c mice (WT) were intraperitoneally infected with *E. granulosus* protoscoleces or injected with the corresponding volume of buffer only. Seven months later, the presence of LL-derived materials in various organs was analysed by histology, using the E492 antibody as a probe **(a)**. Images from representative mice are shown, out of 15 infected and 5 control mice; the bars represent 50 µm. Black arrows indicate KC-like cells staining for LL material; empty arrowheads indicate occasional staining for LL material in spleen and lymph node. The staining observed in adipocytes (upper left-hand corner of the infected mouse mesenteric lymph node section) was also observed in adipocytes present in kidney and lung sections, irrespective of infection, and must therefore represent a cross-reaction of the antibody. No staining was observed in sections probed with the secondary reagent only. The localization of LL materials in KC from infected mice was confirmed by fluorescence histology, using the E492 antibody as probe and either Clec4F or F4/80 as KC markers **(b)** (original magnification x630). The uptake of LL-derived materials by non-parenchymal liver cells of infected BALB/c mice was also analysed by flow cytometry, using PNA as probe. Results are shown for KC, LSEC and non-KC CD11b^+^ cells, in terms of percentages of positive cells **(c)** and mean fluorescence intensities **(d)**, and representative dot-plots for the three cell types are shown in **(e)**. The positive cell threshold was established using staining controls. In addition, WT and Clec4f^-/-^ C57BL/6 mice were similarly infected and the uptake of LL-derived materials analysed by flow cytometry and presented **(f – h)** as for the BALB/c mice. Results in (a – e) are from a single experiment and results in (f - h) are pooled from two independent experiments, shown in blue and red. In (c), (d), (f) and (g), data shown correspond to individual mice and their median values are indicated. The statistical analyses were carried out by the modified Wilcoxon-Mann-Whitney (c, d) and Kruskal-Wallis tests (f, g).

The above results show that in infected animals LL mucins circulate systemically and become selectively concentrated in the liver. Within the liver, the signal for LL mucins was specifically concentrated in KC-like cells (Fig. 7 a). The LL signal in KC-like cells was observed in 12 out of 15 infected mice analysed. The cells taking up LL materials were formally identified as KC by dual-color fluorescence using both F4/80 and Clec4F as KC markers, and antibody E492 as probe for LL materials (Fig. 7 b). Contrary to previous experiments injecting pLL, no signal was detected in hepatocytes. This may be related to lower loads of LL material, as suggested by the lower intensity in the staining in KC in the infection setting in comparison to the pLL injection experiments (compare Fig. 6 e and 7 a).

The selective LL material uptake by KC among the liver non-parenchymal cells was confirmed by flow cytometry (Fig. 7 c – h and S10). Although the data did not reach statistical significance, a sub-population of non-KC CD11b^+^ cells appeared to take up LL materials (Fig. 7 c – d). Also, the frequency of non-KC CD11b^+^ cells within all liver non parenchymal cells was significantly increased by infection, suggesting cell recruitment (Fig. S11 a). The cell surface expression of Clec4F in KC was not altered by infection (Fig. S12 a).

We next analysed infected C57BL/6 WT and Clec4f^-/-^ mice. In these mice, the parasite burdens were low: the median burden in the WT C57BL/6 mice was 0.5 mL (this figure corresponds to the two infection experiments fully analysed, as an additional experiment with very low parasite burdens was excluded from analysis). Although the median parasite burden was 3-fold lower in Clec4f^-/-^ mice than in WT mice, the trend did not reach statistical significance (Fig. S8 b). By flow cytometry, uptake of LL material by KC was detected, in WT but not in Clec4F^-/-^ mice (Fig. 7 f - h). No significant uptake was detected in LSECs (Fig. 7 f, g and S10 b). In non-KC CD11b^+^ cells, there was a trend towards low-level, Clec4F-independent uptake (Fig. 7 f - h), which did not reach significance. The frequency of non-KC CD11b^+^ cells was not significantly altered by infection (Fig. S 11 b). Again, Clec4F surface expression in the KC of WT mice was not altered by infection (Fig. S12 b).

## Discussion

In this article we show that native LL materials shed from *E. granulosus sensu stricto* hydatids are taken up selectively by KC, in a fashion mostly dependent on Clec4F. It is thought that pathogens capable of immune evasion engage innate receptors that normally bind altered self, as opposed to receptors specialized in detecting pathogens and triggering inflammatory responses (42). *E. granulosus* is well adapted for immune evasion (2), and the fact that its most abundant surface molecules engage a receptor involved in the clearing of aged host platelets (20, 21) seems to be in line with the above principle.

KC are specialized in removing unwanted materials from the blood. Our results imply that the LL mucins (which even after their artificial solubilisation behave as having very high molecular sizes (26)) have the capacity to circulate systemically, thus reaching KC from a distant anatomical site. This applies to exogenous LL mucins initially present in the peritoneal cavity in particulate format, which must therefore either circulate as particles and/or partly solubilize *in vivo*. It also applies to the LL materials released by live parasites growing in the peritoneal cavity.

In addition to KC, both LSEC and certain non-KC CD11b^+^ cells were observed to take up LL materials. This was in both cases independent of Clec4F, and in some experimental settings it was enhanced in the absence of Clec4F (Fig. 4 d, 6 d and S6 d). Uptake of materials by non-KC CD11b^+^ cells was apparent after injection of LL mucins in particulate presentation (pLL; Fig. 6 a – d and S6) but not in a soluble presentation (sLL; Fig. 1, 4 and 5). This contrast suggests that the species released *in vivo* from the large LL particles and capable of circulating are (or at least include) small particles, as opposed to fully soluble mucins like those making up sLL. The fact that uptake by the non-KC CD11b^+^ population was clear after injection of a high-dose bolus of pLL whereas it was merely hinted in experimental infection (Fig. 7 c - h) is most probably explained by this uptake mechanism becoming relevant only when the uptake capacity of KC is overwhelmed. It is unlikely that the non-KC CD11b^+^ cell population taking up LL materials derives from peritoneal cavity macrophages, as it is composed of F4/80^-^ cells (Fig. 6 c and S6 c), whereas most resident peritoneal macrophages as well as peritoneal macrophages reported to enter the liver upon injury to this organ are F4/80^hi^ (43). We instead favour the possibility that the population being discussed corresponds to monocytic cells present in the liver or recruited in response to LL particles physically trapped in the liver sinusoids.

Hepatocytes could be an additional target cell for LL materials in the liver, as suggested by the weak but significant staining observed in this cell type after pLL injection (Fig. 6 e). The obvious candidate receptor to mediate this apparent uptake is the ASGR (44). Ligand uptake by hepatocytes via the ASGR is reported to have an upper size limit of 70 nm (45). Thus the signal in hepatocytes observed in our experiments with pLL (which has particle sizes in the micrometer range (13)) may correspond to uptake of soluble mucins released from the particles *in vivo*. In the same experiments as well as in the infection experiments (Fig. 7), the preferential uptake by KC in relation to hepatocytes surely reflects the differential capacity of KC to phagocytose particles (even if additionally, Clec4F may bind LL mucins better than the ASGR, as suggested by *in vitro* evidence (14)). Our results do not allow to weigh the importance of hepatocytes in the capture of fully soluble injected LL materials (sLL).

Part of the uptake of LL materials by KC was observed to be Clec4F-independent (Fig. 5 a and 6 a, b). Clec4F-independent uptake by KC, as well as by non-KC CD11b^+^ cells and LSEC as previously mentioned, may be mediated by natural anti-carbohydrate antibodies that bind the LL mucins, either directly or via complement activation (46). In the context of chronic infection, the possibilities for Clec4F-independent uptake are even broader than after LL mucin injection. In the first place, the mucins will be opsonized by specific (homologous) antibodies (7, 46). In second place, contrary to the artificially prepared (soluble and insoluble) mucin preparations, LL materials shed from live parasites must contain the major non-mucin structural component of the LL, i.e. calcium inositol hexakisphosphate (calcium Ins*P*_6_) (47–49). Calcium Ins*P*_6_ binds host C1q and initiates limited activation of complement (46, 50). In fact, the observed uptake of LL materials by phagocytes at the experimental infection site (Fig. 7 a) must also be Clec4F-independent, as peritoneal macrophages are not expected to express Clec4F (17). Given the ample possibilities of Clec4F-independent capture, it is noteworthy the Clec4F nonetheless has a clearly detectable role in LL material uptake (by KC), especially in the infection context.

As mentioned, in human infections affecting solid organs, LL debris accumulate in the parasite’s vicinity (11). Our results show the presence of LL materials at the experimental infection site but also as mentioned the capture of these materials by the liver (Fig. 7 a). This implies that the accumulation of LL debris in the parasite’s vicinity is moderated by the circulation and capture by the liver of materials derived from this debris. This is important because, although moderate amounts of LL particles probably have immune modulatory effects that favour the parasite, large amounts of LL particles accumulating at the infection site would probably foster inflammation (12, 13, 51, 52). Inflammation might also be brought about the LL materials remaining in circulation and possibly being deposited in the glomeruli or other sites. Indeed, if as we speculated above, monocytes are recruited to the liver in response to trapped LL particles, then Clec4F-dependent KC capture may be important to avoid liver inflammation. This aspect may not have been fully captured in our flow cytometry and histology experiments because of liver perfusion prior to analysis.

The events downstream of LL material capture by KC are certainly worth studying. KC can present antigens to T cells present in the vascular space, and this normally leads to tolerogenic responses that can have a systemic impact (23, 53, 54). The analysis of immune responses in WT and Clec4f^-/-^ mice infected with *E. granulosus* will be the subject of a future communication.

Ancestrally, the larval stages of the parasites belonging to the taeniid family (including those of the genus *Echinococcus*) probably infected the liver of rodent hosts (6, 55). In this context, the interaction of materials shed by the parasites with a receptor (Clec4F) specifically expressed by phagocytic cells present in the same organ (KC) may have been evolutionarily advantageous. This may have been so because LL material uptake by KC would have avoided particle build-up and consequent inflammation in the liver sinusoids and/or other sites, and/or because KC are poised for tolerogenic antigen presentation. The extant species *E. multilocularis* dwells almost exclusively in the liver of rodents (besides accidentally infecting humans). The LL of this species also interacts with Clec4F, as suggested by experiments using recombinant Clec4F (14) and by the presence in this species’ LL of the blood antigen P_1_ motif (56). Thus, host Clec4F is likely an important player in *E. multilocularis* infection.

The natural hosts of larval *E. granulosus sensu lato* are mostly ungulates, including sheep, cattle, pigs, goats, camels and horses. Like in rodents, in ungulates Clec4F also appears to be expressed in the liver (16). Therefore, the uptake of LL materials by KC via Clec4F probably takes place also in the ungulate natural hosts. However, in ungulates Clec4F appears to be additionally expressed in secondary lymphoid organs (16), opening the possibility that non-KC, anatomically more widely distributed, phagocytes may act as additional sinks for LL materials. Cystic echinococcosis in ungulate hosts, although mostly caused by *E. granulosus sensu stricto*, is also caused by *E. ortleppi*, *E. canadensis*, and *E. equinus* (depending on the host species) (3). The blood antigen P_1_ motif, shown in this work to be a major target of Clec4F binding to the LL, is in all likelihood abundant in the LL of all *E. granulosus sensu lato* species, as it is abundant in the LL of *E. multilocularis* (56). Therefore, although our data were obtained using *E. granulosus sensu stricto*, Clec4F is very probably relevant to cystic echinococcosis in ungulates irrespective of the infecting species within the *E. granulosus sensu lato* cluster.

As mentioned, humans do not express Clec4F with a functional lectin domain (16). Humans are normally a dead-end for this parasite’s life-cycle, meaning that human *Echinococcus* infection has not been shaped through evolution. Although *E. multilocularis* infection in humans is usually accompanied by strong local inflammation (and tissue destruction), this is not clearly the case with *E. granulosus* (10, 57). It is a possibility that for *E. granulosus*, which grows more slowly than *E. multilocularis*, Clec4F-independent clearance mechanisms are in most infection settings sufficient to avoid excessive build-up of LL debris. It is conceivable that such Clec4F-independent mechanisms in humans include uptake via lectins expressed in KC with specificities related to Clec4F. As mentioned, the uptake of de-sialylated platelets by mouse KC requires Clec4F as well as MGLs or ASGR (20, 21). It seems likely that a mechanism for de-sialylated platelet uptake by KC based on a set of collaborating lectins exists throughout mammals, even if in humans such a system does not include Clec4F. Our previous screening using recombinant receptors yielded inconclusive results with respect to the capacity of human MGL and ASGR to interact with the LL mucins (14). When probing synthetic sugars, human ASGR and MGL appeared to bind less well than Clec4F to the P_1_ motif (after normalizing by binding to a known target of all 3 receptors) (14). However, given the sheer abundance and significant breadth of Gal-based motifs in the LL, it cannot be ruled out that ASGR and/or MGL may be able to internalize LL materials in the absence of Clec4F. We plan to determine if KC in liver samples from humans infected with *E. granulosus*, as well as with *E. multilocularis*, stain for LL materials using the probes introduced in this work. We also posit that the characterization of the anatomical distribution, mechanisms of cell uptake, and possible inflammatory effects of LL materials in Clec4f^-/-^ mice will provide useful insight on the biology of human cystic echinococcosis.

## Statement of Ethics

Mouse studies followed the ARRIVE guidelines. Experimental protocols were individually reviewed and approved by the Institutional Animal Ethics Committees of the Institut Pasteur Montevideo or the Universidad de la República, Uruguay, as detailed in the Materials and Methods section.

## Acknowledgements

The authors are indebted to Marcela Cucher (University of Buenos Aires, Buenos Aires, Argentina) for advice with *Echinococcus* genotyping and to Gustavo Salinas (Universidad de la República and Institut Pasteur de Montevideo, Uruguay) for helpful discussions.

‘For the purpose of open access, the authors have applied a Creative Commons Attribution (CC BY) licence to any Author Accepted Manuscript version arising’.

## Author contributions

AB: conceptualization, investigation. CM, LS, JN, MS: investigation. GS, MNM, APM, MC: methodology. TLH, SLH, GME: resources. CG: formal analysis (statistics). TFEB, CC: conceptualization, **s**upervision. SJJ: conceptualization, funding acquisition, supervision, writing – review & editing. AD: conceptualization, funding acquisition, project administration, supervision, writing – original draft preparation.

## Supplementary Figures

**Fig. S1. Summary of preparation of LL materials.** The preparation of pLL and sLL (periodate- or mock-treated) is summarized in **(a)**. Periodate oxidation of terminal monosaccharides and further monosaccharides carrying *cis*-diols generates reactive aldehyde groups, which are reduced to alcohols using sodium borohydride. The mock treatment consists of the borohydride step only. The effectiveness of the periodate treatment was verified in terms of abrogation of PNA binding by to the mucins in a dot-blot format **(b)**. Similar incorporation of the biotin tag to periodate- and mock-treated sLL was verified using streptavidin-peroxidase in dot-blot format **(c)**.

**Fig. S2. Gating strategy for liver cells.** Gating strategies are shown for KC, LSEC, and non-KC CD11b^+^ cells. For non-KC CD11b^+^ cells, which tend to be substantially smaller than KC, it was necessary to include an additional doublet exclusion step at the end of the gating strategy.

**Fig. S3 (related to** Fig. 3**). Reactivity of PNA, E492 antibody and recombinant Clec4F with liver non-parenchymal cells of control mice.** Non-parenchymal cells from WT C57BL/6 mice were stained for flow cytometry with PNA, E492 antibody or recombinant mouse Clec4F, followed by appropriate secondary probes. The staining controls correspond to samples stained in the same way as the positive ones except that the primary probe was omitted.

**Fig. S4 (related to** Figs. 4 and 5**). Representative flow plots for liver non-KC CD11b^+^cells.** sLL was injected in WT and Clec4f^-/-^ C57BL/6 mice, and liver non parenchymal cells analysed after 20 min **(a)** or 22 h **(b)**, as described in Figs. 4 and 5 respectively. Representative flow plots are shown for non-KC CD11b^+^ cells, which were not found to take up sLL significantly.

**Fig. S5 (related to** Fig. 5**). Surface expression of Clec4F does not change upon injection of sLL.** WT C57BL/6 mice were injected with sLL or vehicle only (PBS) as described in Fig. 5, and 22 h later KC were analysed for surface expression of Clec4F by flow cytometry. Data shown, in terms of geometric means of fluorescence intensity (MFI), arise from 3 independent experiments, shown in different colors. The data were normalized over the median value of mice injected with PBS only in each experiment.

**Fig. S6 (related to** Fig. 6**). Uptake of pLL by liver non-parenchymal cells measured by flow cytometry using antibody E492.** WT and Clec4f^-/-^ C57BL/6 mice were injected with pLL and material uptake was analysed as in Figure 6 except that the E492 antibody was used as probe. Results are shown in terms of percentages of positive cells **(a)** and mean fluorescence intensities **(b)**, and representative dot-plots are shown in **(c)**. An estimation of the fraction of sLL taken up by each of the cell populations under study, calculated as explained in the Materials and Methods section, is shown in **(d)**; the statistics shown compare the fractions taken up by KC. The positive cell threshold was established using staining controls. The data arise from two independent experiments, represented in blue and red in (a-c) and indicated as 1, 2 in (d). In (a-c), individual mice and their median values, and in (d) only medians, are shown. The statistical analyses were carried out by the Mack-Skillings test.

**Fig. S7 (related to** Fig. 6**). Frequency of non-KC CD11b^+^ cells in mice injected with pLL.** WT and Clec4f^-/-^ C57BL/6 mice were injected with pLL or vehicle only (PBS) as described in Fig. 6, and 22 h later the frequency of non-KC CD11b^+^ cells within all live cells was measured by flow cytometry. Data arise from 2 independent experiments, shown in different colors.

**Fig. S8 (related to** Fig. 6**). Surface expression of Clec4F does not change upon injection of pLL.** WT C57BL/6 mice were injected with pLL or vehicle only (PBS) as described in Fig. 6, and 22 h later KC were analysed for surface expression of Clec4F by flow cytometry. Data shown, in terms of geometric means of fluorescence intensity (MFI), arise from 2 independent experiments, shown in different colors. The data were normalized over the median value of mice injected with PBS only in each experiment.

**Fig. S9 (related to** Fig. 7**). Parasite burdens of mice infected with *E. granulosus*.** WT BALB/c mice **(a)** or WT and Clec4f^-/-^ C57BL/6 mice **(b)** were infected with *E. granulosus* as described in Fig. 7. Parasite burdens are expressed in terms of total volumes of metacestodes, measured by water displacement. Results for BALB/c mice correspond to a single experiment and results for C57BL/6 are pooled from 2 experiments, represented in different colours. Individual mice and their overall median values are shown. Parasite burdens were not found to be significantly different between WT and Clec4f^-/-^ C57BL/6 mice by the Kruskall-Wallis test.

**Fig. S10 (related to** Fig. 7**). Representative flow plots for LSEC in infections.** WT BALB/c mice **(a)** or WT and Clec4f^-/-^ C57BL/6 mice **(b)** were infected with *E. granulosus* and liver non-parenchymal cells analysed by flow cytometry as described in Fig. 7. Representative flow plots are shown for LSEC, which were not found to take up LL materials.

**Fig. S11. Frequencies of non-KC CD11b^+^ cells in *E. granulosus* experimental infections.** WT BALB/c mice **(a)** or WT and Clec4f^-/-^ C57BL/6 mice **(b)** were infected with *E. granulosus* and liver non-parenchymal cells analysed by flow cytometry as described in Fig. 7. Frequencies of non-KC CD11b^+^ cells within total live cells are plotted. The statistical analysis was carried out by the modified Wilcoxon-Mann-Whitney method.

**Fig. S12. Experimental *E. granulosus* infection does not alter the expression of Clec4F in KC.** WT BALB/c mice **(a)** or WT C57BL/6 mice **(b)** were infected with *E. granulosus* as described in Fig. 7. KC were analysed for surface expression of Clec4F by flow cytometry. Data shown, in terms of geometric means of fluorescence intensity (MFI), arise from 1 experiment for each of the mouse strains shown. The data were normalized over the median value of the control mice in each experiment.

