## Supplemental Table and Figures for "Mucins shed from the laminated layer in cystic echinococcosis are captured by Kupffer cells via the lectin receptor Clec4F"

**Barrios *et al*. submitted.**

**Supplementary Table 1. Probes used in flow cytometry.** All extracellular (pre-permeabilization) probes were diluted in PBS containing 0.1% w/v BSA and 2 mM EDTA. All intracellular (post-permeabilization) probes were diluted in permeabilization buffer.

| **Probe name** | **Dilution or concentration** | **Clone** | **Manufacturer** | **Secondary reagent (if relevant)** | **Used before or after permeabilization** |
|---|---|---|---|---|---|
| LIVE/DEAD™ fixable far red dead cell stain kit | 1/500 | Not applicable | Thermo (Invitrogen) | | before |
| Anti-CD45 APC-Cy7 | 1/200 | 30-F11 | Biolegend | | before |
| Anti-F4/80 PE-Cy7 | 1/300 | BM8 | Biolegend | | before |
| Anti-CD11b PerCP-Cy5 | 1/400 | M1/70 | Biolegend | | before |
| Anti-CD31 Brilliant Violet 421 | 1/200 | 390 | Biolegend | | before |
| Anti-Clec4F | 1/50 | Goat polyclonal | R&D | donkey-anti-goat Alexa Fluor 488 | before (except when indicated otherwise) |
| Anti-goat IgG Alexa Fluor 488 | 1/700 | Donkey polyclonal | Thermo (Invitrogen) | | As for anti-Clec4F |
| Anti-CD16/32 | 1/2000 | 2.4G2 | Biolegend | | before |
| E492-biotin | 0.6 µg/mL | E492 (mouse $IgG_3$ monoclonal) | Prepared in house | Streptavidin-PE | after |
| PNA-biotin | 1/1500 | Not applicable | Vector Laboratories | Streptavidin-PE | after (except when indicated otherwise) |
| Streptavidin PE (used as primary probe to | 1/200 | Not applicable | eBioscience | | after (except when indicated otherwise, in |

| | | | | | |
|---|---|---|---|---|---|
| detect sLL-biotin, or used following PNA-biotin or E492-biotin) | | | | | an experiment in combination with PNA-biotin) |

**Figure S1**

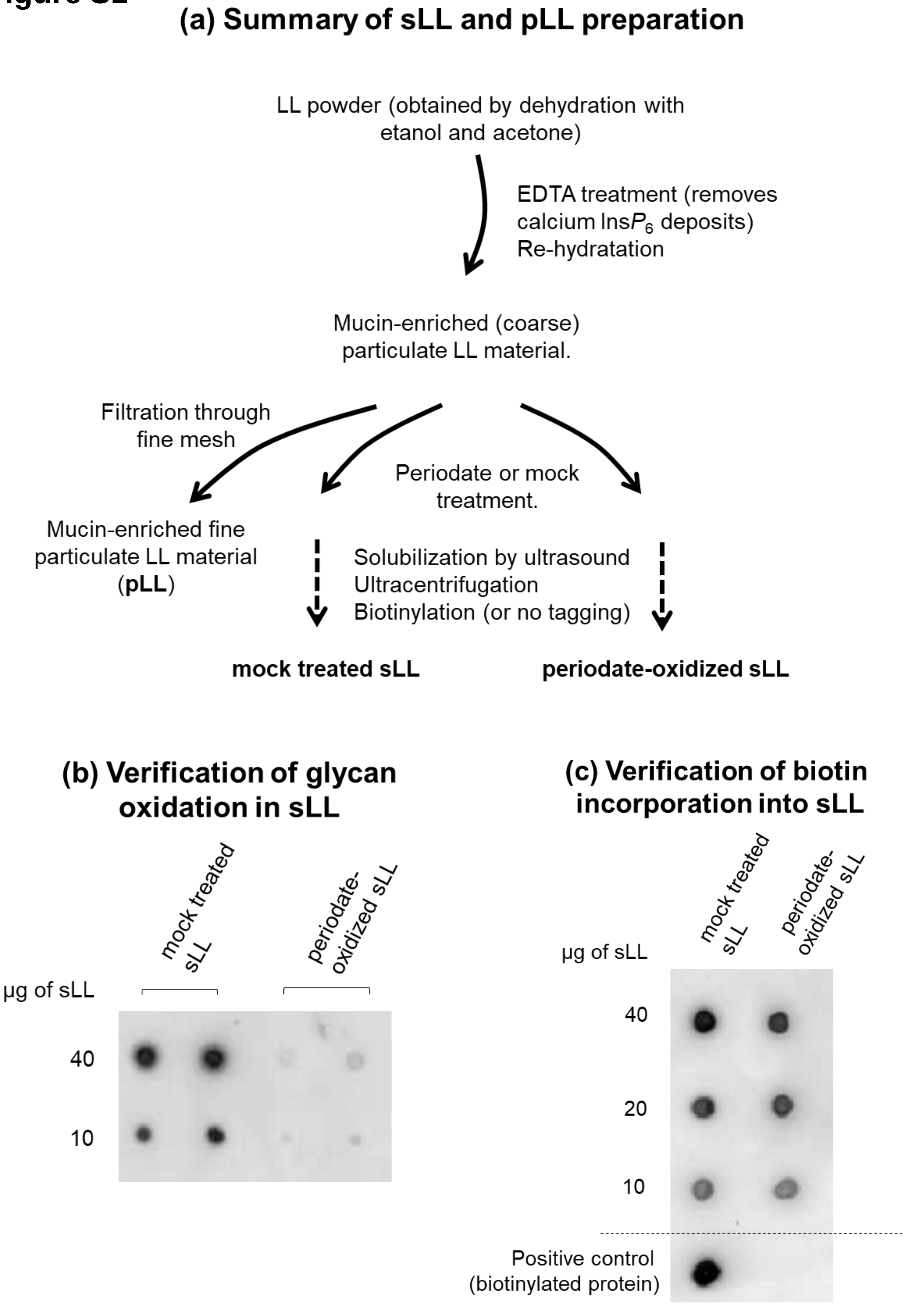
(a) Summary of sLL and pLL preparation
LL powder (obtained by dehydration with etanol and acetone)
EDTA treatment (removes calcium Ins$P_6$ deposits)
Re-hydratation
Mucin-enriched (coarse) particulate LL material.
Filtration through fine mesh
Periodate or mock treatment.
Mucin-enriched fine particulate LL material (pLL)
Solubilization by ultrasound
Ultracentrifugation
Biotinylation (or no tagging)
mock treated sLL
periodate-oxidized sLL
(b) Verification of glycan oxidation in sLL
mock treated sLL
periodate-oxidized sLL
µg of sLL
40
10
(c) Verification of biotin incorporation into sLL
mock treated sLL
periodate-oxidized sLL
µg of sLL
40
20
10
Positive control (biotinylated protein)

### Figure S2

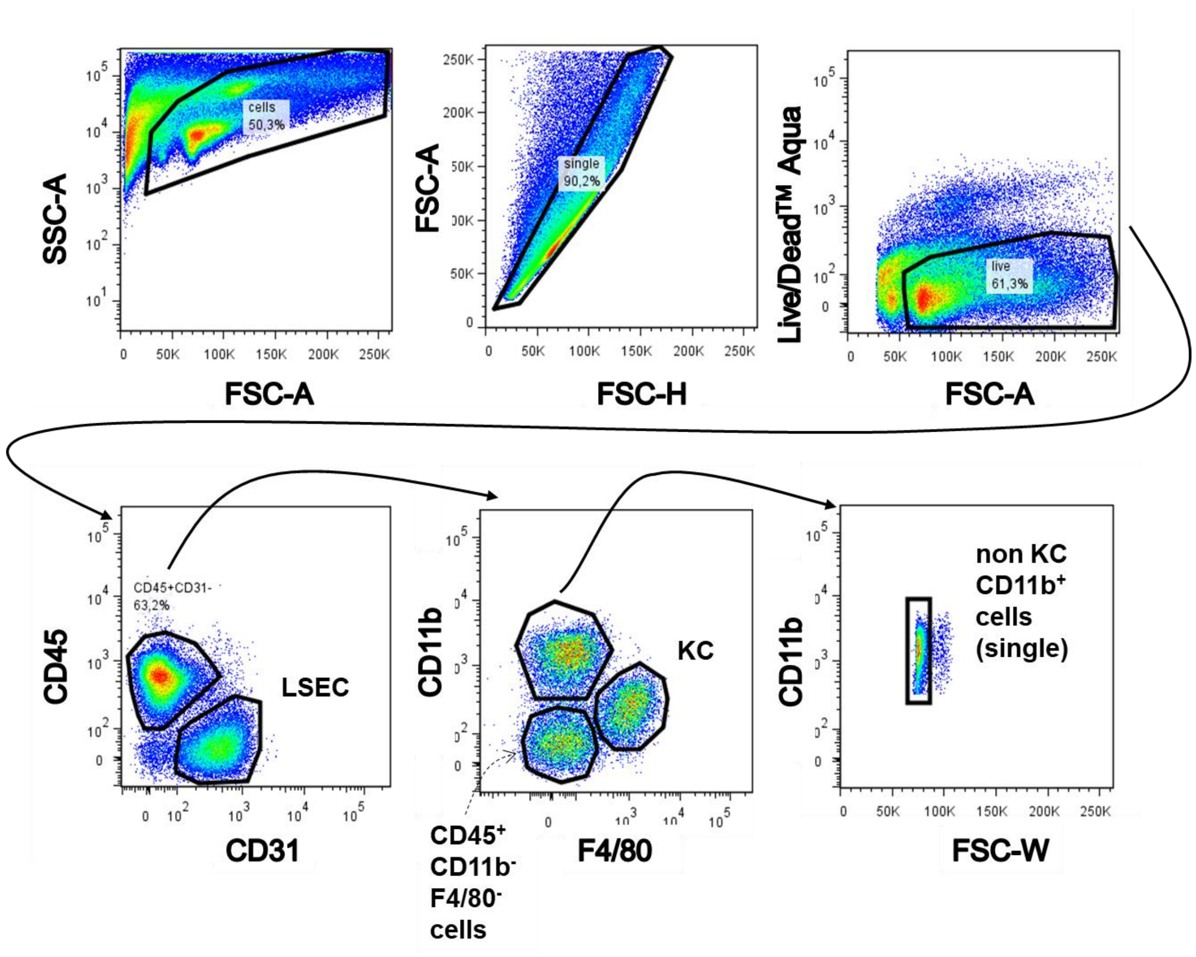

SSC-A
FSC-A
cells
50,3%
FSC-A
FSC-H
single
90,2%
Live/Dead™ Aqua
FSC-A
live
61,3%
CD45
CD31
CD45+CD31-
63,2%
LSEC
CD11b
F4/80
KC
CD45⁺
CD11b⁻
F4/80⁻
cells
CD11b
FSC-W
non KC
CD11b⁺
cells
(single)

### Figure S3

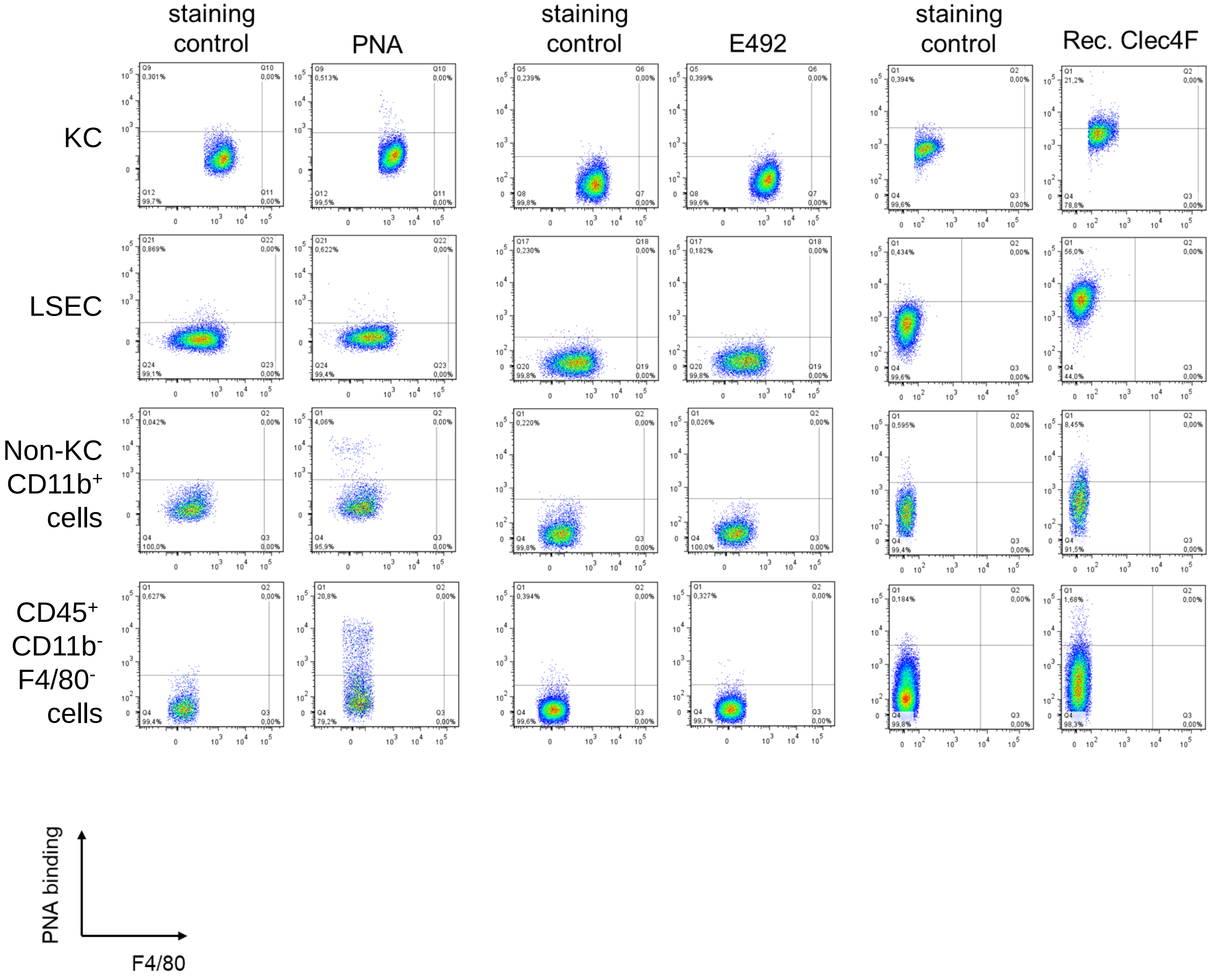
staining control
PNA
staining control
E492
staining control
Rec. Clec4F
KC
LSEC
Non-KC CD11b+ cells
CD45+ CD11b- F4/80- cells
PNA binding
F4/80

### Figure S4

#### (a) sLL injection and analysis after 20 min

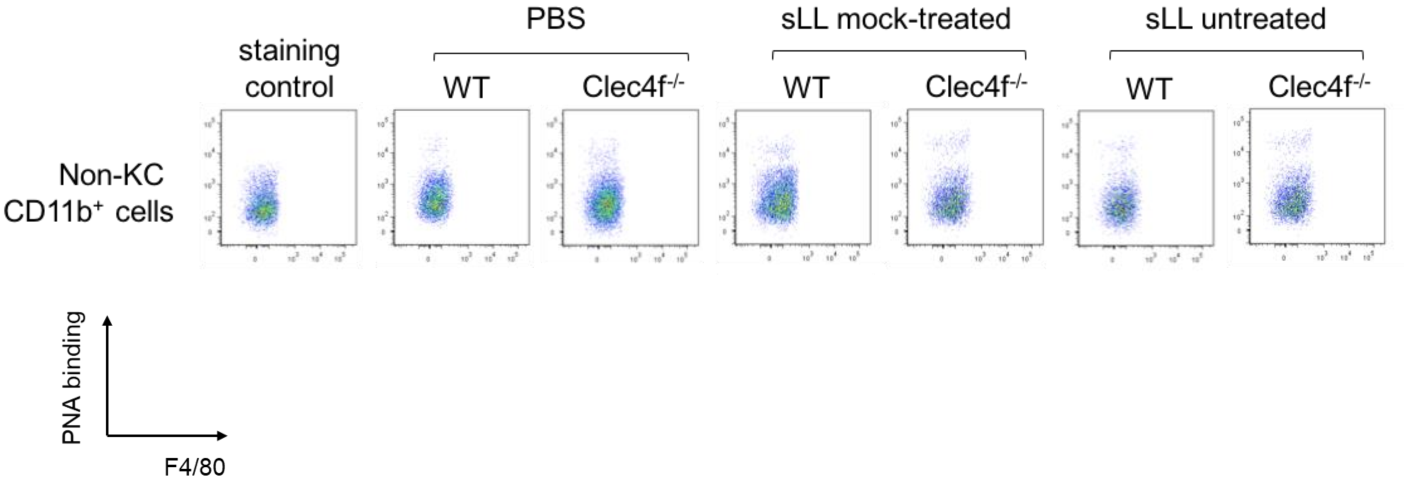


#### (b) sLL injection and analysis after 22 h

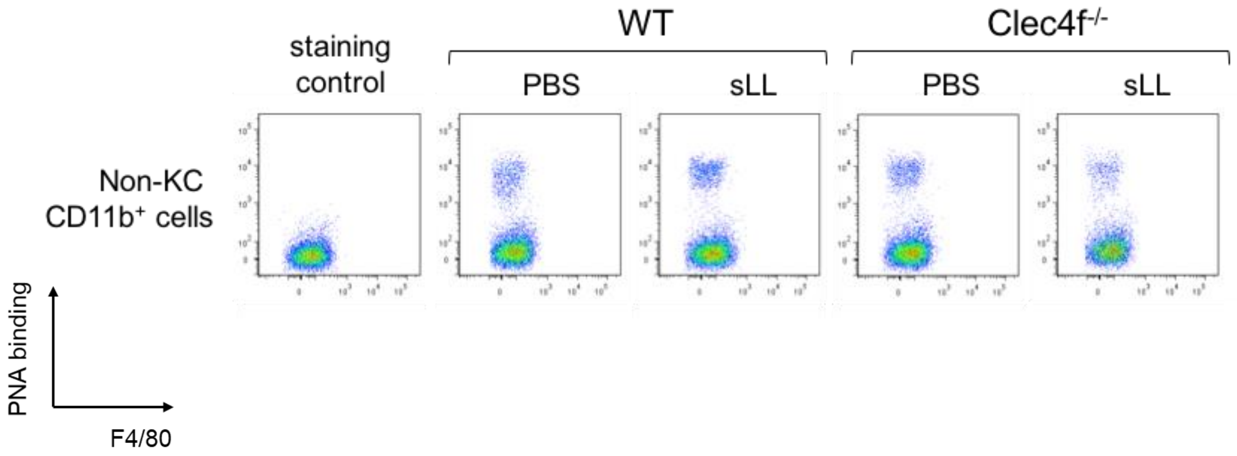

### Figure S5

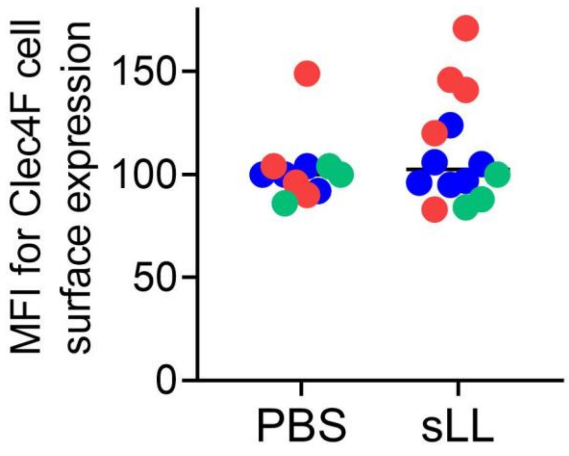

### Figure S6

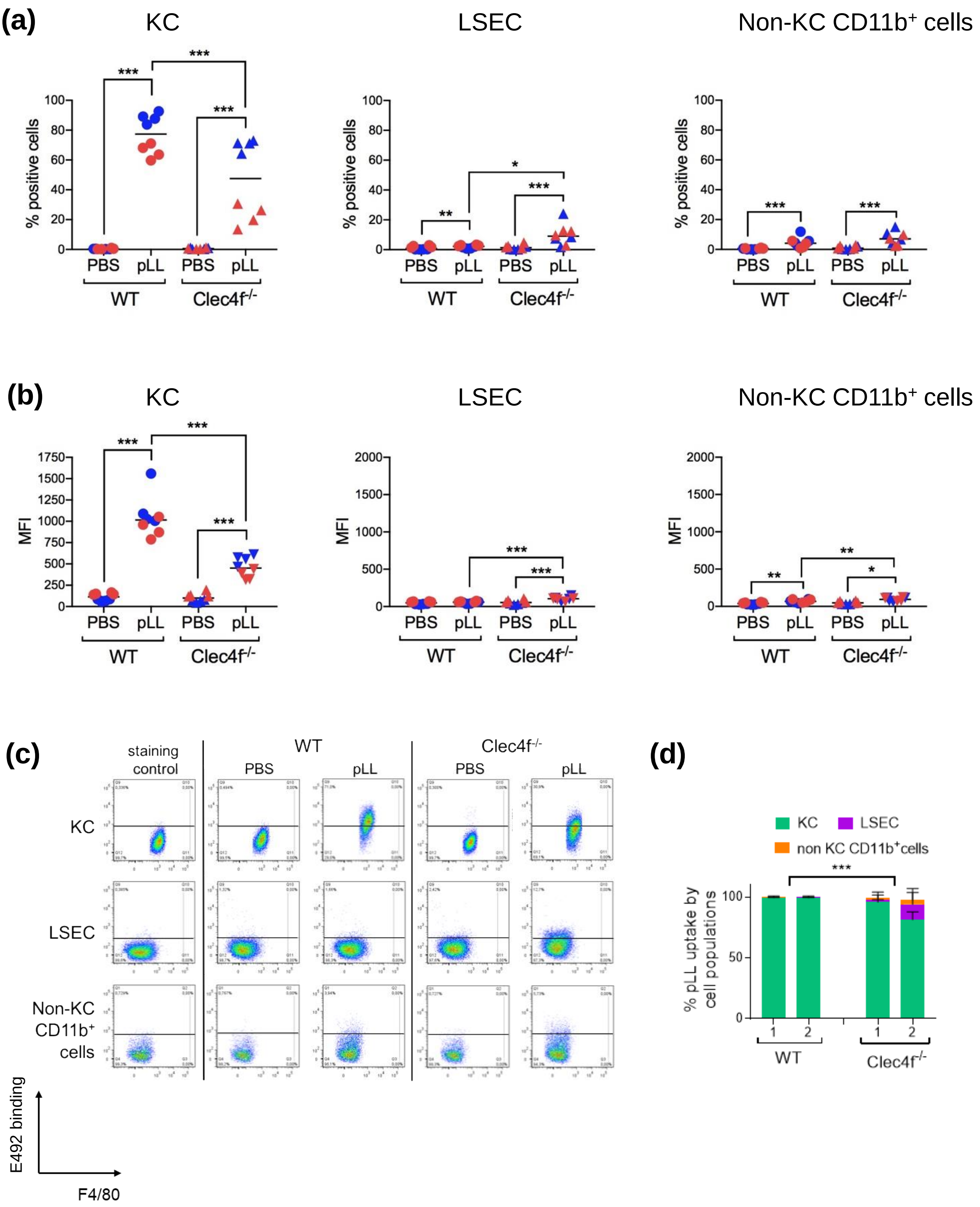
(a)
KC
LSEC
Non-KC CD11b+ cells
% positive cells
PBS
pLL
WT
Clec4f-/-
(b)
MFI
(c)
staining control
KC
LSEC
Non-KC CD11b+ cells
E492 binding
F4/80
(d)
KC
LSEC
non KC CD11b+cells
% pLL uptake by cell populations

**Figure S7**

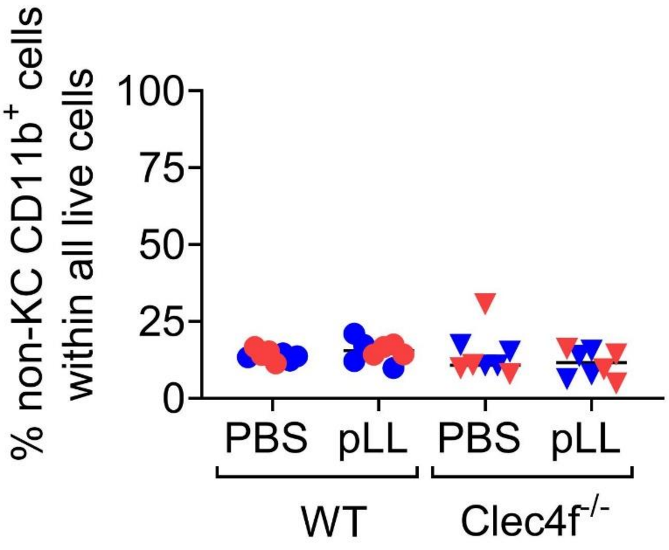

% non-KC CD11b+ cells within all live cells
100
75
50
25
0
PBS
pLL
PBS
pLL
WT
Clec4f-/-

**Figure S8**

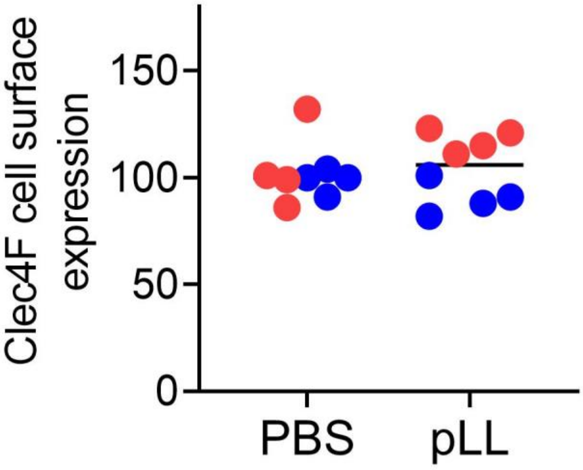

### Figure S9

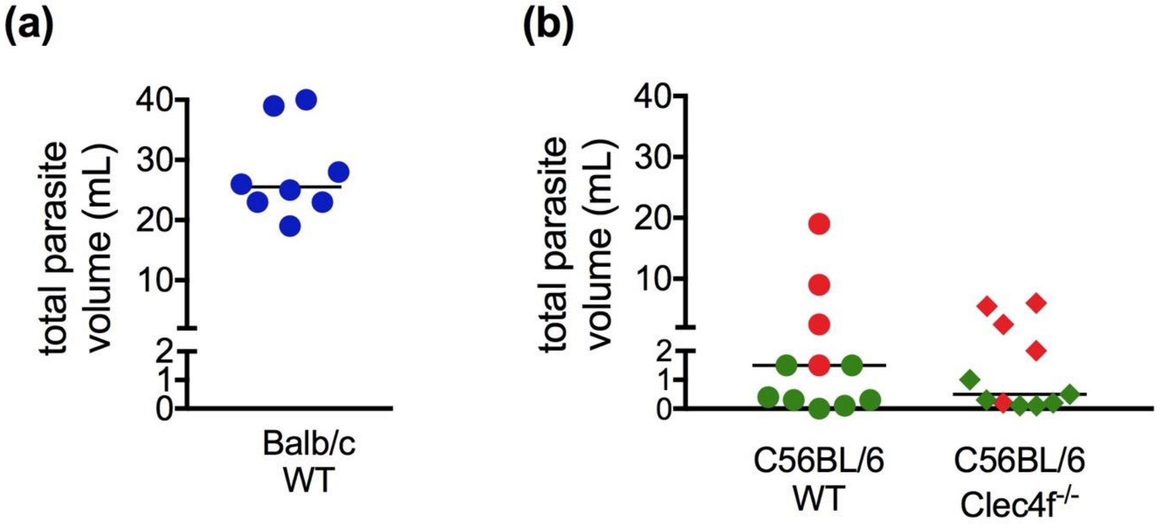
(a)
total parasite volume (mL)
40
30
20
10
2
1
0
Balb/c WT
(b)
total parasite volume (mL)
40
30
20
10
2
1
0
C56BL/6 WT
C56BL/6 Clec4f-/-

### Figure S10

**(a)** Infection in Balb/c mice

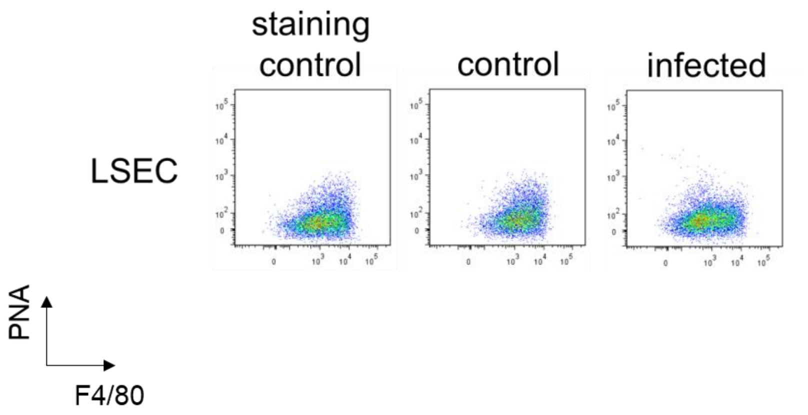


**(b)** Infections in WT and Clec4f$^{-/-}$ C57BL/6 mice

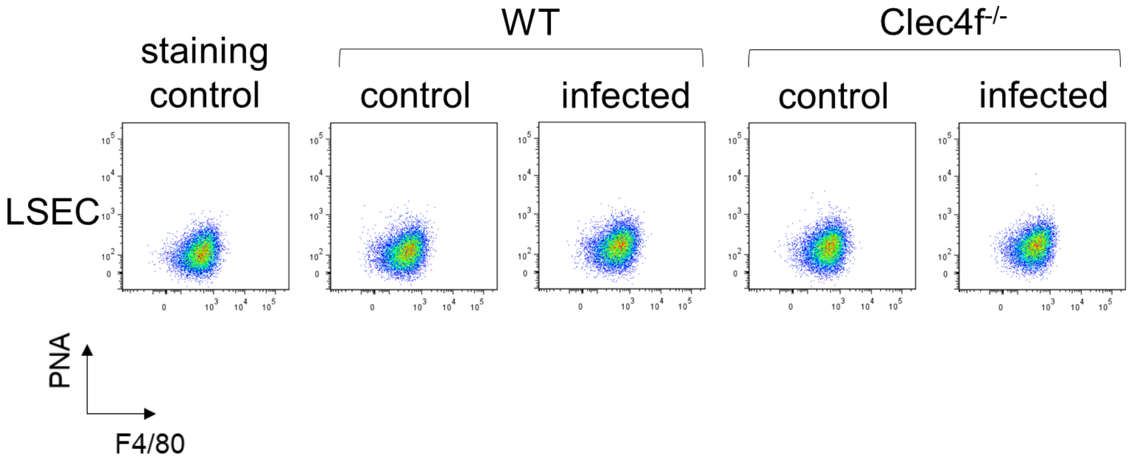

### Figure S11

**(a)** Infection in Balb/c mice

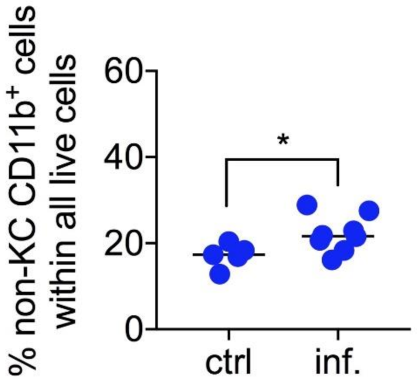


**(b)** Infections in WT and Clec4f$^{-/-}$ C57BL/6 mice

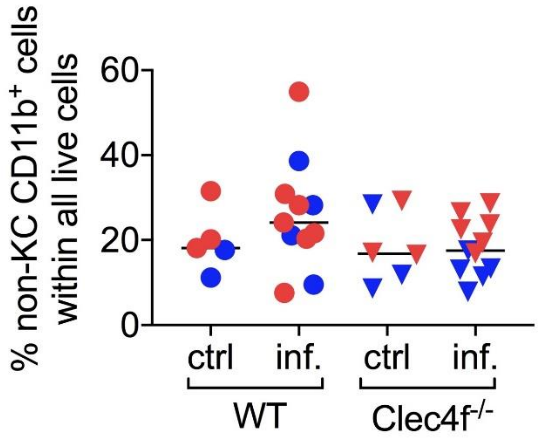

### Figure S12

**(a)** Infection in Balb/c mice

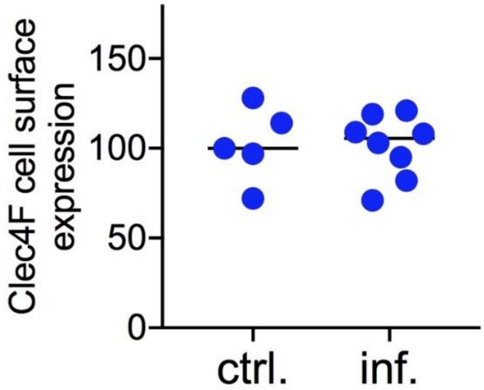


**(b)** Infections in WT and Clec4f$^{-/-}$ C57BL/6 mice

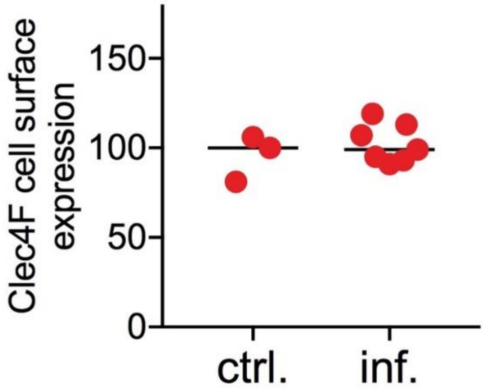
